# CyclinD2-mediated regulation of neurogenic output from the retinal ciliary margin is perturbed in albinism

**DOI:** 10.1101/2022.01.27.477954

**Authors:** Nefeli Slavi, Revathi Balasubramanian, Melissa A. Lee, Michael Liapin, Rachel Oaks-Leaf, John Peregrin, Anna Potenski, Carol Troy, M. Elizabeth Ross, Eloisa Herrera, Stylianos Kosmidis, Simon W. M. John, Carol A. Mason

**Author notes:** Authors contributed equally to this work (Nefeli Slavi, Revathi Balasubramanian). Corresponding author: Carol Mason. Co-corresponding author: Nefeli Slavi. Competing interests: the authors declare no competing interests.

## Abstract

In albinism, aberrations in the ipsi-/contralateral retinal ganglion cell (RGC) ratio compromise the functional integrity of the binocular circuit. We focus here on the mouse ciliary margin zone (CMZ), a neurogenic niche at the embryonic peripheral retina, to investigate developmental processes regulating RGC neurogenesis and identity acquisition. We found that the mouse ventral CMZ has the competence to generate predominantly ipsilaterally-projecting RGCs, but this competence is altered in the albino visual system due to CyclinD2 downregulation and disturbed temporal control of the cell cycle. Consequently, albino as well as CyclinD2-deficient pigmented mice exhibit a diminished ipsilateral retinogeniculate projection and compromised depth perception. Pharmacological stimulation of calcium channels in albino mice, known to upregulate CyclinD2 in other cell types, augmented CyclinD2-dependent neurogenesis of ipsilateral RGCs, and improved stereopsis. Together, these results implicate CMZ neurogenesis and its regulators as critical for the formation and function of the mammalian binocular circuit.

**Highlights:**

1. The mouse ventral CMZ produces predominantly ipsilateral RGCs.
2. In the albino visual system, CyclinD2 downregulation leads to delayed G1/S transition toward mitotic exit of CMZ progenitors.
3. Perturbations in the temporal control of cell cycle by CyclinD2 lead to reduced Zic2^+^ RGCs and consequently, a diminished ipsilateral retinogeniculate projection and compromised depth perception.
4. Calcium channel modulation during embryogenesis normalizes the levels of CyclinD2 and restores binocular vision in albino mice.

## Introduction

A key evolutionary feature of the mammalian visual system is a higher degree of binocular overlap, attributed to a change in the eye position from the sides to the front of the head. To properly form the binocular circuit, specific numbers of retinal ganglion cells (RGCs) from each retina project axons to either the ipsilateral or the contralateral brain hemisphere after partial decussation at the optic chiasm (Murcia-Belmonte and Erskine, 2019; Petros et al., 2008). The resultant convergence and precise integration of visual information from both eyes to binocularly-driven cortical neurons constitute the foundation of stereopsis (Cumming and DeAngelis, 2001; Howarth et al., 2014; Scholl et al., 2013). While genes specific to ipsi- and contralateral RGCs are being uncovered (Kuwajima et al., 2017; Lo Giudice et al., 2019; Morenilla-Palao et al., 2020; Shekhar et al., 2021; Su et al., 2021; Wang et al., 2016), a complete mechanism that links early regulatory processes driving the production of ipsi- and contralateral RGCs to specific aspects of binocular vision has not been described.

Classic birthdating experiments show that RGC neurogenesis begins principally at the center of the retina at the junction between the optic cup and the optic stalk, and spreads as a wave toward the periphery (Young, 1985). In fish, amphibian tadpoles, and birds, retinal neurons can also be generated at the peripheral-most region of the embryonic retina, the ciliary margin zone (CMZ), facilitating eye growth and regeneration (Fischer and Reh, 2000; Kubota et al., 2002; Wan et al., 2016). Recent studies describe a combinatorial code of growth factors driving CMZ vs neural retina specification in the vertebrate eye cup (Balasubramanian et al., 2021), and report that the mouse CMZ also contains a population of RGC progenitors that have molecular identity distinct from progenitor cells in the neural retina (Belanger et al., 2017; Marcucci et al., 2016; Tropepe et al., 2000). However, crucial questions regarding the neurogenic properties of the mammalian CMZ remain unresolved. First, are there CMZ-specific mechanisms that control RGC fate acquisition and projection path to the brain? Second, is RGC neurogenesis from the CMZ critical to the formation of the binocular circuit, thereby supporting depth perception?

In humans with albinism, a genetic disorder affecting melanin biosynthesis, stereoscopic vision is severely impaired. Developmentally, albinism is characterized by delayed center-to-periphery retinal neurogenesis (Bhansali et al., 2014), and diminished expression of Zic2, the key transcription factor conveying ipsilateral RGC fate (Garcia-Frigola et al., 2008; Herrera et al., 2003; Morenilla-Palao et al., 2020), which results in an abnormally small ipsilateral eye-to-brain projection (Herrera et al., 2003) and aberrant targeting of recipient regions in the dorsal lateral geniculate nucleus (dLGN) (Rebsam et al., 2012). Ipsilateral RGCs are positioned adjacent to the CMZ. Consequently, we asked whether and how the capacity of the CMZ to generate ipsilateral RGCs is affected in the albino mouse, and interrogate its impact on the functional integrity of the binocular visual system.

Here we demonstrate that the mouse ventral CMZ is a rich source of ipsilateral RGCs. Within the CMZ, we focus on the temporal control of the cell cycle, which determines the timing of neurogenesis thereby influencing neuronal identity during differentiation (Cavalieri et al., 2021; Cepko et al., 1996; Marcucci et al., 2018; Osterhout et al., 2014; Pilaz et al., 2016b; Rossi et al., 2017; Tripodi et al., 2011). Unlike the neural retina, the CMZ is enriched in CyclinD2, a cell cycle protein that regulates the proliferation and fate of radial glial cells and intermediate progenitors in cortical and subcortical neurogenic zones (Glickstein et al., 2009; Petros et al., 2015; Pilaz et al., 2016a; Tsunekawa et al., 2012; Tsunekawa et al., 2014). We show that expression of CyclinD2 in CMZ progenitors is required for appropriate G1-to-S phase transition, securing their mitotic exit in time to express Zic2 and acquire an ipsilateral RGC fate. In contrast to the pigmented eye, we find that in the albino, downregulation of CyclinD2 leads to an asynchronous cell cycle and reduced RGC neurogenesis from the CMZ during a time window critical for Zic2 expression. Ultimately, both albino and conditional CyclinD2-deficient pigmented mice exhibit depth perception deficits during a visually-guided behavioral task due to disproportionate binocular inputs to the dLGN. Lastly, we demonstrate that pharmacological modulation of CyclinD2 expression in the CMZ increases ipsilateral RGCs, prevents axon misrouting in the albino binocular circuit, and restores depth perception.

## Results

### Ipsilateral RGC neurogenesis is prominent in the pigmented CMZ but is reduced in the albino

Cells of the CMZ expressing the transcription factor Msx1 have been described as a source of retinal neurons during mouse development (Belanger et al., 2017; Marcucci et al., 2016). To characterize the output of CMZ neurogenesis, we used Msx1 CreERT2; tdTomato pigmented and albino mice to fate map progenitors of the ventral CMZ, and examined RGC production in the time window critical for ipsilateral RGC fate acquisition (embryonic days E13.5-E15.5) (Herrera et al., 2003; Marcucci et al., 2018) (Fig. 1A-A’). At each of the three time-points (E13.5, 14.5, 15.5) of Msx1 CreERT2 activation, the number of Msx1^+^ cells was similar between pigmented and albino ventral CMZ (Fig. 1B-B’).

**Figure 1:**
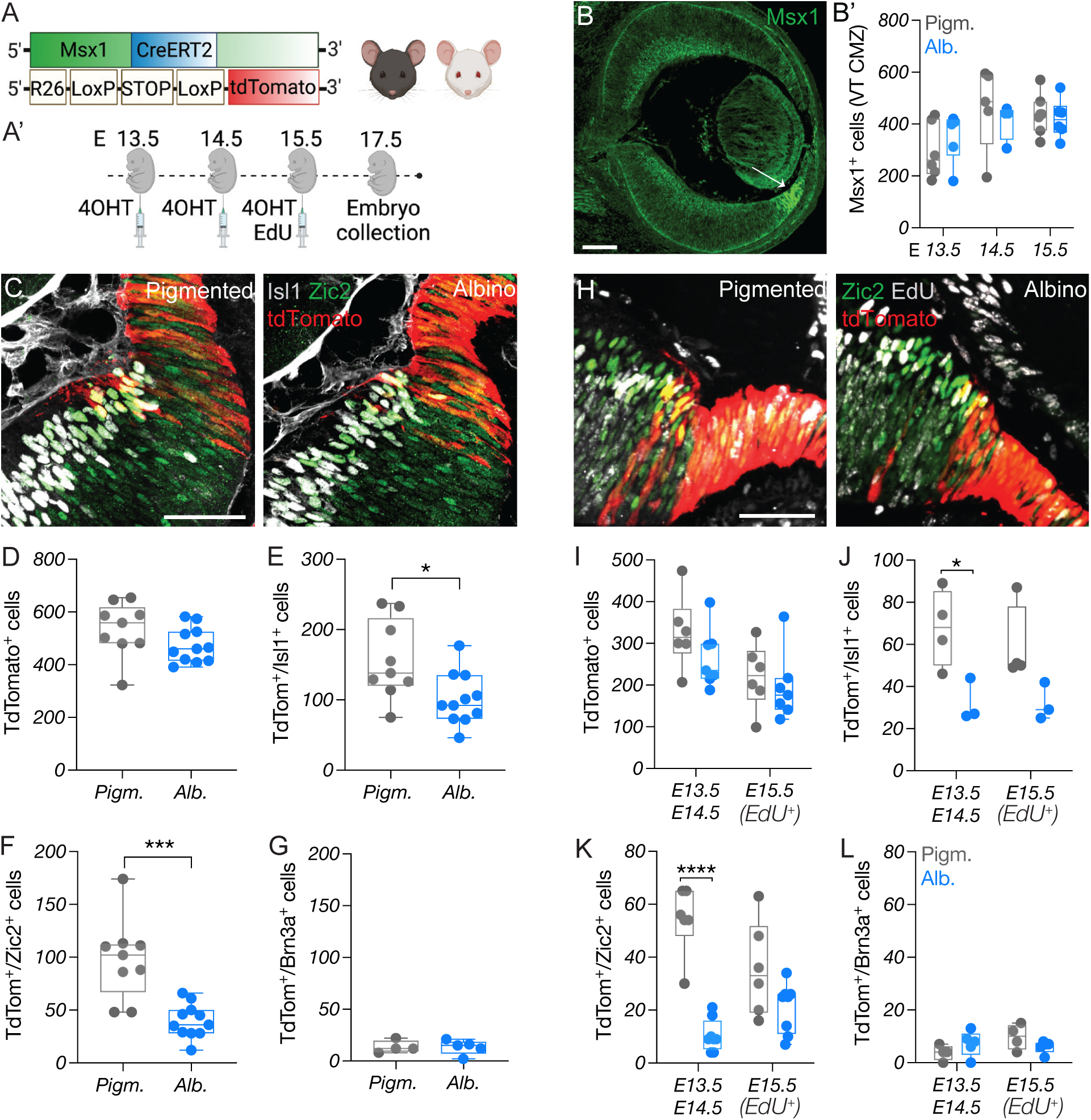
Ipsilateral RGC neurogenesis is prominent in the pigmented CMZ but is reduced in the albino. A) Schema of the elements used to generate Msx1 CreERT2; tdTomato; Tyr mice. A’) Timeline of 4-OHT and EdU injections and sample collection. B) Example immunostaining of Msx1 at E14.5, showing its exclusive expression in the CMZ (arrow). Scale bar: 100μm. B’) Quantification of Msx1^+^ cells in the ventral CMZ of pigmented and albino mice at E13.5, E14.5 and E15.5. C) Immunostaining of tdTomato, Islet1, and Zic2 in the pigmented and albino ventrotemporal CMZ and retinal crescent at E17.5. Scale bar: 50μm. D-G) Quantification of tdTomato^+^ (D), tdTomato^+^/Islet1^+^ (E), tdTomato^+^/Zic2^+^ (F) and tdTomato^+^/Brn3a^+^ (G) cells in the ventrotemporal retina of pigmented and albino mice at E17.5. H) Immunostaining of tdTomato, EdU, and Zic2 in the pigmented and albino ventrotemporal retina and retinal crescent E17.5. Scale bar: 50μm. I-L) Quantification of tdTomato^+^ (I), tdTomato^+^/Islet1^+^ (J), tdTomato^+^/Zic2^+^ (K) and tdTomato^+^/Brn3a^+^ (L) cells born at E13.5-14.5 (EdU^-^) or E15.5 (EdU^+^) in the ventrotemporal retina of pigmented and albino mice at E17.5.

Upon tamoxifen administration, and by E17.5, the Msx1^+^ domain of the CMZ was well-labeled with tdTomato, while some tdTomato^+^ cells were localized in the adjacent neural retina, in agreement with previous studies (Belanger et al., 2017; Marcucci et al., 2016) (Fig. 1C). Neurogenesis from the CMZ increased gradually along the rostro-caudal axis of the eye, with more tdTomato^+^ cells found in the temporal segment of the retina (Fig. S1A). This pattern was detected in both pigmented and albino mice, and the total number of CMZ-derived cells found in the neural retina across both genotypes was non-distinguishable (Fig. 1D). To detect and quantify postmitotic RGCs that derived from the CMZ, we examined Islet1 expression, marking all differentiated RGCs. Again, we observed that RGC neurogenesis from the CMZ was elevated in the temporal compared with the nasal retina of pigmented and albino mice (Fig. S1B). The albino retina, however, contained fewer tdTomato^+^/Islet1^+^ RGCs (Fig. 1E), suggesting a compromised capacity for RGC neurogenesis of the albino ventral CMZ.

Next, we asked whether curtailed RGC neurogenesis in the albino CMZ impacts the ipsilateral (Zic2^+^), the contralateral (Brn3a^+^) RGC population, or both. We especially focused on the ventrotemporal quadrant of the retina, a locus where ipsilateral RGCs are generated exclusively from E13.5 to E16.5, followed by late-born contralateral RGCs from E15.5 until birth. We found that the CMZ of both genotypes indeed produced ipsilateral RGCs. However, the generation of Zic2^+^ RGCs from the albino CMZ was reduced compared to the pigmented CMZ (Fig. 1C, F and Fig. S1C). This reduction of Zic2^+^ RGC neurogenesis from the albino CMZ resembled the reduction of Zic2^+^ RGC generation from center-to-periphery neurogenesis (Zic2^+^/tdTomato^-^ cells) in the albino ventrotemporal retina (Fig. S1E). In contrast to Zic2^+^/tdTomato^+^ RGCs, extremely few Brn3a^+^/tdTomato^+^ cells were detected in both the pigmented and albino retina, indicating that the majority of RGCs derived from the ventral CMZ are ipsilateral (Fig. 1G and Fig. S1D), and suggesting that until E17.5 contralateral RGCs originate instead from the neural retina. No differences were noted in the number of Brn3a^+^/tdTomato^-^ ventrotemporal RGCs between pigmented and albino retina (Fig. S1F).

To probe for differences in the pace of neurogenesis between the pigmented and albino CMZ, that might be responsible for the reduction in CMZ-derived Zic2^+^ RGCs in the albino, we combined fate mapping with birthdating by injecting EdU at E15.5 (Fig. 1A, 1H). This allowed us to distinguish tdTomato^+^ cells born at E13.5 and E14.5 (EdU^-^) from those born at or after E15.5 (EdU^+^). The CMZ of pigmented mice contributed tdTomato^+^ cells (Fig. 1I), including Zic2^+^ RGCs (Fig. 1J-K), to the neural retina throughout this time window at a gradually decreasing rate, in line with the typical maturation of CMZ. In contrast, the albino CMZ produced fewer Zic2^+^ RGCs at E13.5-E14.5, compared to pigmented CMZ (Fig. 1J-K). At all time points (E13.5-E15.5), very few contralateral (Brn3a^+^) RGCs were born in both the pigmented and albino CMZ (Fig. 1L).

Collectively, our findings show that the ventral CMZ produces RGCs with a primarily ipsilateral fate, and that reduced RGC neurogenesis in the albino CMZ between E13.5 and E14.5 partially accounts for the diminished ipsilateral RGCs population that marks albinism.

### scRNA-Seq in the pigmented and albino CMZ

To investigate the early processes that govern neurogenesis from the CMZ and are perturbed in the albino visual system, we conducted scRNA-Seq on FACS-isolated αCre; tdTomato^+^ cells from E13.5 mouse peripheral retinae (Fig. 2A-B). The use of αCre; tdTomato^+^ mice allowed for enrichment in CMZ cells, as upon activation of the retina-specific regulatory element of Pax6 α-enhancer, the retinal periphery including the CMZ is fluorescently labeled (Marquardt et al., 2001). We sequenced 6,089 pigmented and 3,676 albino cells at a mean depth of 3,200 genes per cell. Unsupervised clustering and UMAP representation of the developing peripheral retina (Fig. 2C-D and Fig. S2A) revealed eleven cell clusters (Balasubramanian et al., 2021; Clark et al., 2019; Lo Giudice et al., 2019). First, a group of progenitor cells (positive for *Sfrp2, Hes1, Fgf15*) expressed markers specific to the CMZ (cluster 1) (*Ccnd2, Wfdc1, Msx1, Gja1*). Adjacent to the CMZ cluster, three groups of progenitor cells (RPC1-3; clusters 0, 2, 9) (positive for *Sfrp2, Ccnd1, Hes1, Fgf15*) could be distinguished from each other based on distinct patterns of cell cycle gene expression (e.g., *Hist1h2ap, Hist1h2ae, Cenpf, Ube2c, Prc1, Nusap1*) (Fig. S2B). The RPC clusters were connected to a young neurogenic cell population (cluster 3) expressing *Atoh7, Hes6*, and *Neurog2*, as well as cell cycle genes (H1 histone family genes, *Top2a*), followed by a mid- (*Atoh7, Hes6)* and a more mature neurogenic cluster (clusters 10, 5), enriched in *Atoh7* and *Hes6*, as well as in RGC, horizontal and amacrine cell genes. Emanating from the neurogenic cells, and opposite to the CMZ cluster, we observed two separate branches, one with young and old RGCs (clusters 4, 7) (*Isl1, Pou4f2*-enriched), and the other that included a horizontal/amacrine cell cluster (cluster 8) (*Ptf1a, Lhx1, Onecut1/2*-enriched) as well as a photoreceptor precursors/cones cluster (cluster 11) (*Otx1, Neurod1/4, Crx*-enriched).

**Figure 2:**
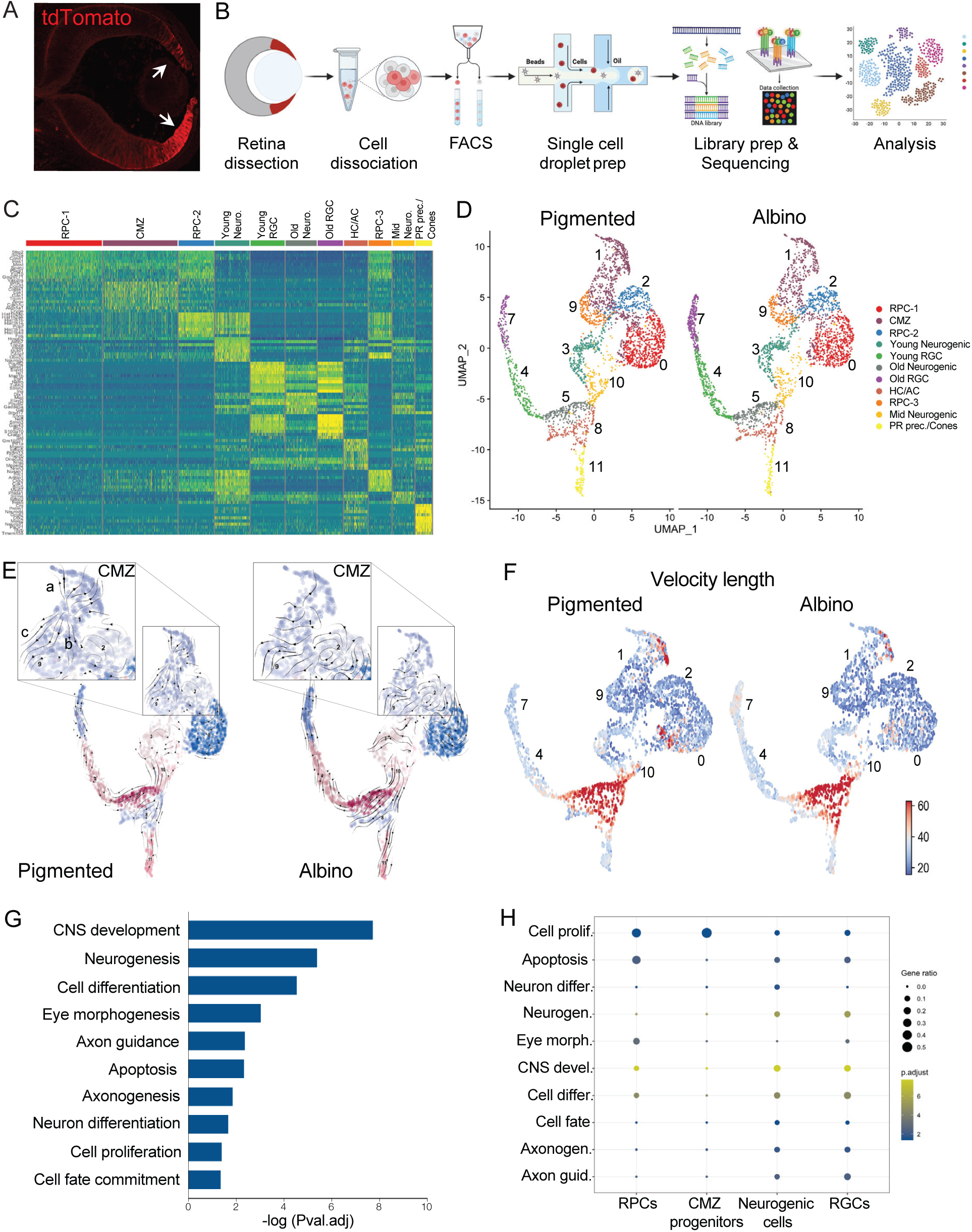
scRNA-Seq in the pigmented and albino CMZ. A) Example image of tdTomato expression (arrows) in the CMZ and peripheral retina of αCre; tdTomato mice at E13.5. B) Schema summarizing the scRNA-Seq work flow. C) Heat map of differential gene expression in single-cell clusters. D) UMAP representation of single-cell clusters in the pigmented and albino datasets. E) RNA Velocity vector stream projected on UMAP of single-cell clusters in the pigmented and albino datasets. F) Length of velocity vector quantifying arrow length, embedded on UMAP of single-cell clusters in the pigmented and albino datasets. G) GO terms corresponding to all differentially expressed genes between pigmented and albino datasets. H) GO term (from G) enrichment across cell states. RPC: retinal progenitor cell, CMZ: ciliary margin zone cell, Neuro: neurogenic cell, RGC: retinal ganglion cell, HC: horizontal cell, AC: amacrine cell, PR: photoreceptor

We next investigated changes to the rate and direction of cell state transitions using RNA velocity analysis (Fig. 2E-F, Fig. S3A). Cells within the CMZ cluster of the pigmented eye (cluster 1) had three distinct predicted trajectories (Fig. 2E), which reflected transition towards: a) terminal ciliary margin cells indicated by arrows moving upwards towards the tip of cluster 1 (CMZ), b) self-renewing RPCs indicated by arrows moving towards clusters 2 and 9 (RPC-2, RPC-3), or c) neurogenic states as indicated by arrows exiting towards cluster 7 (RGCs). Velocity across all three predicted trajectories was altered in cells of the albino CMZ, such that the arrows went in varied directions (Fig. 2E), potentially reflecting a disruption in the progression of differentiation. Specifically, the velocity of cells transitioning into mature ciliary margin cells (e.g., expressing *Mitf, Wls*) was reduced as seen by the quantification of vector length (Fig. 2F, Fig. S3A). The albino RPC-1 population (cluster 0) also displayed slower velocities of cell state progression towards neurogenic fate (cluster 10) compared to the pigmented RPC-1 population (cluster 1) (Fig. 2F, Fig. S3A). Conversely, the velocity of retinal cells differentiating towards RGCs (cluster 7) was slightly higher in the albino, presumably caused by an increase in contralateral RGC fate (Fig. 2F, Fig. S3A). Together, these data suggest that neurogenesis from the albino CMZ does not follow the directionality and tempo of the pigmented CMZ at E13.5.

Lastly, we performed a gene ontology (GO) analysis of genes that were differentially expressed (DE) between pigmented and albino, focusing on the CMZ, RPC, neurogenic, and RGC clusters (Fig. 2G). The GO term “ regulation of cell proliferation” (DE genes *Jun, Ccnd2, Adam10, Ptn, Hmgcr, Hes5, Gnai2)* was the major difference between the pigmented and albino CMZ and retinal progenitors, supporting the alterations in RNA velocity length (Fig. 2H, Fig. S3B). The GO terms of differentiation, neurogenesis, and cell fate commitment (*Neurog2, Atoh7, Ascl1, Dlx1, Gap43, Islr2, Srrm4, Tenm3, Cadm1, Mdk, Dpysl2, Dcx, Ina, Casp3*) were enriched in neurogenic cells, whereas DE genes characterizing RGC neurogenesis, axonogensis, and axon development (*Cxcr4, Top2b, Cntn2, Mapt, Pak3, Nefl, Nefm*) were enriched in RGCs (Fig. 2H and Fig. S3C-F), in line with the transcriptional changes reflecting cell state transitions during retinal development. Lastly, genes associated with ipsilateral RGC fate (e.g., *Zic2, Igfbp5, Pcsk2*) (Herrera et al., 2003; Lo Giudice et al., 2019; Wang et al., 2016) were downregulated in the albino neurogenic and RGC clusters (Fig. S3D), corroborating our fate mapping results, whereas genes linked to contralateral fate (e.g., *Pou3f1, Nrp1, Fgf12*) were increased (Fig. S3E).

### Cell cycle is perturbed in the albino CMZ

Studies on the albino retina have associated lack of pigment with spatiotemporal defects in retinal neurogenesis, such as mitotic spindle orientation, and delayed pace of center-to-periphery cell production and maturation (Bhansali et al., 2014; Cayouette et al., 2001; Ilia and Jeffery, 1996; Jeffery, 1998; Rachel et al., 2002; Tibber et al., 2006). It is not clear, however, whether regulation of specific cell cycle phases is an important determinant of RGC fate specification, especially in the CMZ.

The finding that the expression of genes regulating cell proliferation was altered in the albino CMZ prompted us to compare the pigmented and albino scRNA-Seq datasets based on S and G2/M phase scores (Fig. 3A). Cell cycle scoring revealed a reduction in the number of cells in both S and G2/M phases in the albino compared to pigmented peripheral retina (Fig. 3B). To validate this observation and to further probe for differences in the progression of pigmented and albino CMZ cells through the cell cycle, we performed dual-pulse birthdating at E13.5 and E14.5 using BrdU and EdU. To identify cells within the cell cycle we additionally stained for Ki67. We counted cells in G1 phase based on the expression of Ki67 and absence of BrdU and EdU labeling (Ki67^+^ only), as well as cells in S phase based on Ki67 and EdU labeling with or without BrdU (Ki67^+^/EdU^+^/BrdU^±^) (Fig. 3C). The albino CMZ contained a greater number of cells in G1 phase compared to the pigmented CMZ, and fewer cells in the S phase (Fig. 3E). We observed no significant differences in the total number of Ki67^+^ CMZ cells between the two groups (Fig. 3D). Next, we quantified the number of CMZ cells in M phase using the mitotic marker PH3 (Fig. 3F), and found fewer mitotic cells in the albino CMZ compared to pigmented at E13.5 and E14.5 (Fig. 3G). Therefore, progenitors in the albino CMZ progress through the cell cycle and reach mitotic exit at a slower pace (Fig. 3H) due to a delay in their transition from G1 to S phase.

**Figure 3:**
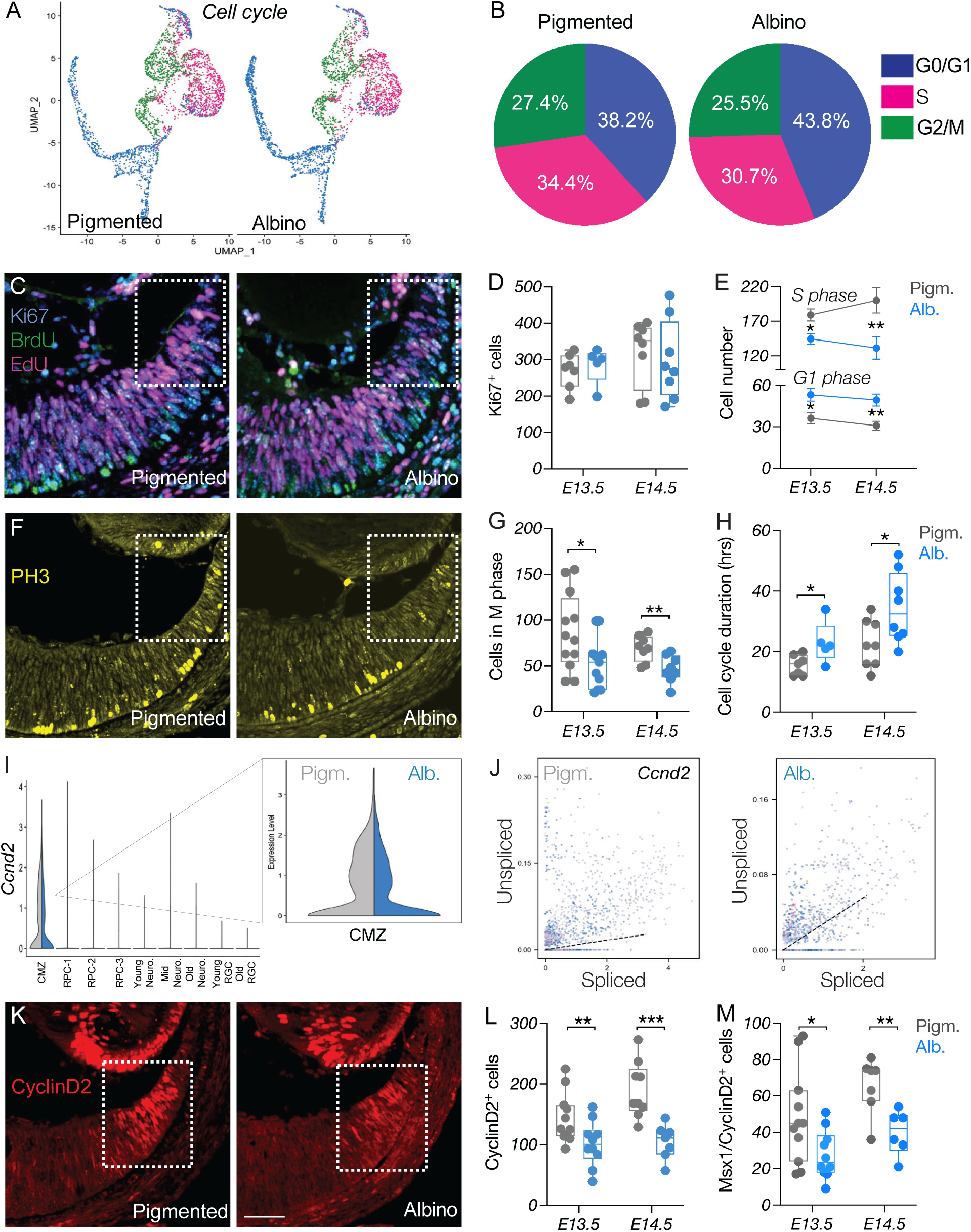
Cell cycle is perturbed in the albino CMZ. A) UMAP representation of single cells based on cell cycle phase. B) Quantification of cells in the G0-G1, S or G2/M phase of the cell cycle, in pigmented and albino datasets. C) Dual-pulse birthdating: Immunostaining of Ki67, BrdU, and EdU in the pigmented and albino ventrotemporal CMZ at E14.5. D) Quantification of Ki67^+^ cells at E13.5 and E14.5. E) Quantification of cells in the G1 and S phases of the cell cycle at E13.5 and E14.5. F) Immunostaining of the mitotic marker PH3 in the pigmented and albino ventrotemporal CMZ at E14.5. G) Quantification of cells in the M phase of the cell cycle (PH3^+^) at E13.5 and E14.5. H) Quantification of cell cycle duration (hours) at E13.5 and E14.5. I) Violin plot showing downregulation of *Ccnd2* in the albino CMZ cell cluster. J) Transcriptional dynamics of *Ccnd2* as a ratio of unspliced to spliced transcripts in the pigmented and albino peripheral retina. K) Immunostaining of CyclinD2 in the pigmented and albino ventrotemporal CMZ at E14.5. Scale bar: 100μm. L-M) Quantification of CyclinD2^+^ (L) and CyclinD2^+^/Msx1^+^ (M) cells at E13.5 and E14.5.

We next sought to pinpoint the cause of this delay. Among the candidates we identified in our scRNA-Seq, the predominant DE gene in the CMZ cluster was *Ccnd2* (Fig. S3B, Fig. 3I), which encodes for CyclinD2, a cell cycle-regulatory protein that has been associated with RGC neurogenesis and identity (Marcucci et al., 2018; Wang et al., 2016). Moreover, cells of the pigmented peripheral retina had a positive *Ccnd2* velocity profile that indicates transcriptional activation, as the production of unspliced *Ccnd2* mRNA exceeded the degradation of its spliced counterpart. In the albino, however, a greater number of cells had negative *Ccnd2* velocity profile, indicating its transcriptional downregulation (Fig. 3J). Consistent with our scRNA-Seq findings, the albino CMZ contained fewer CyclinD2^+^ cells compared to pigmented at E13.5 and E14.5 (Fig. 3K-L). CyclinD2 expression was also reduced in the albino Msx1^+^ CMZ progenitor pool (Fig. 3M).

These results demonstrate that weak CyclinD2 expression in retinal progenitors within the albino CMZ is correlated with disrupted transitioning from the G1 to S phase of the cell cycle, which subsequently impacts the transition to M phase.

### CyclinD2 is a critical regulator of ipsilateral RGC neurogenesis

We hypothesized that delayed cell cycle G1/S progression at E13.5 and E14.5 due to insufficient CyclinD2 expression prevents CMZ cells from exiting the cycle in time to express Zic2, in turn, to establish the proper proportion of ipsilaterally-projecting RGCs. To determine a causal role of CyclinD2 in the temporal regulation of the cell cycle and ipsilateral RGC neurogenesis in the CMZ, we compared pigmented CyclinD2^flox^ (CyclinD2^WT^) mice with pigmented αCre; CyclinD2^flox^ mice lacking CyclinD2 specifically in the CMZ (CyclinD2^cKO^) (Fig. S4A). First, we repeated our dual pulse labeling experiment (Fig. 4A). Similar to the albino CMZ, the CyclinD2^cKO^ CMZ had more cells in G1 and fewer in S phase compared to CyclinD2^WT^ (Fig. 4B). Furthermore, in the CyclinD2^cKO^ CMZ we detected a reduction of PH3^+^ cells (Fig. 4C-D), suggesting that decrease in CyclinD2 expression in the CMZ is associated with a prolonged G1-to-S transition, preventing cells from exiting the cell cycle, as in the albino retina (Fig. 3C-E).

**Figure 4:**
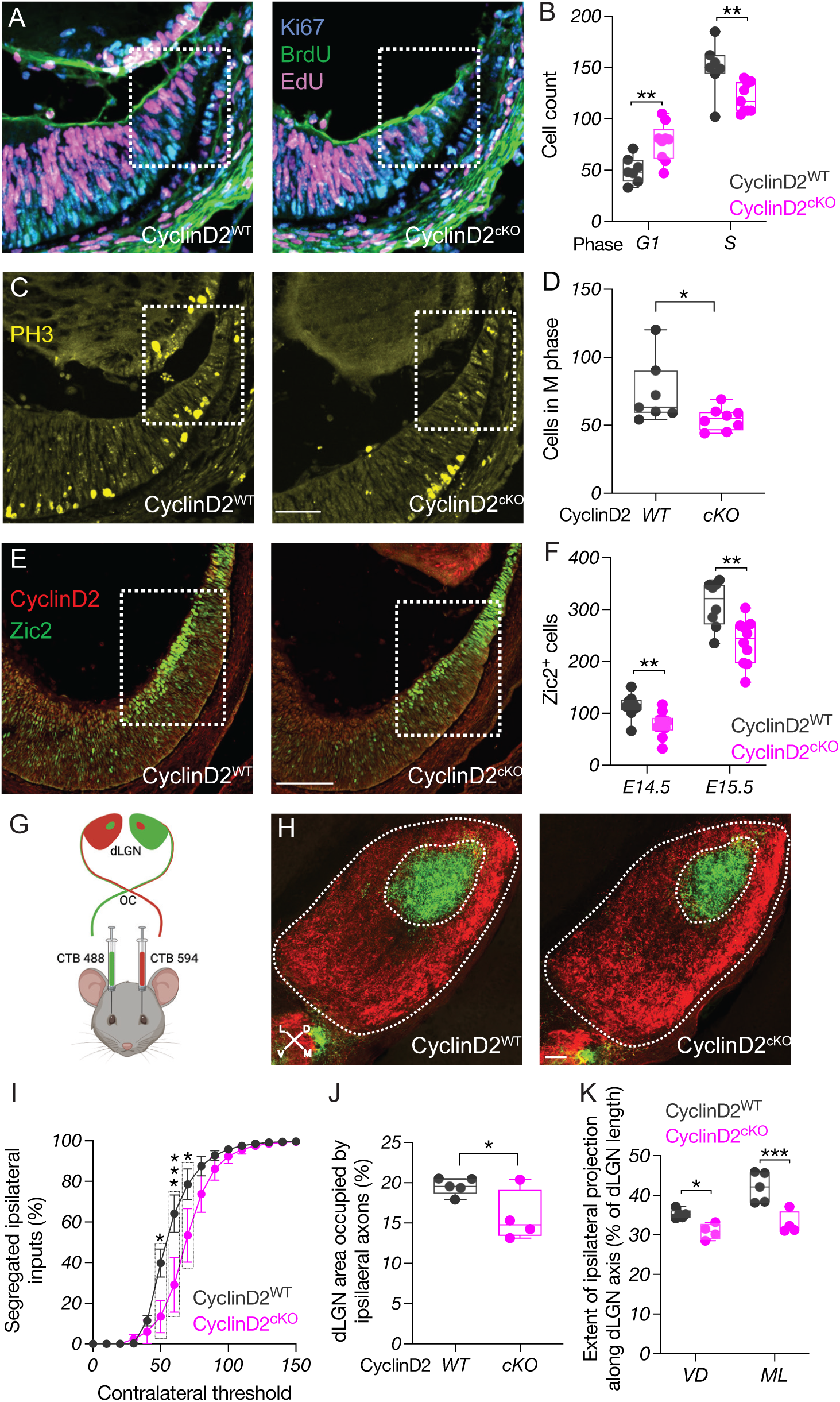
CyclinD2 is a critical regulator of ipsilateral RGC neurogenesis. A) Dual-pulse birthdating: Immunostaining of Ki67, BrdU, and EdU in the CyclinD2^WT^ and CyclinD2^cKO^ ventrotemporal CMZ at E14.5. B) Quantification of cells in the G1 and S phases of the cell cycle at E14.5. C) Immunostaining of the mitotic marker PH3 in the CyclinD2^WT^ and CyclinD2^cKO^ ventrotemporal CMZ at E14.5. Scale bar: 100μm. D) Quantification of cells in the M phase of the cell cycle (PH3^+^) at E14.5. E) Immunostaining of CyclinD2 and Zic2 in the CyclinD2^WT^ and CyclinD2^cKO^ ventrotemporal retina and CMZ at E15.5. Scale bar: 100μm. F) Quantification of Zic2^+^ cells at E14.5 and E15.5. G) Schema of the strategy used for RGC axon tracing to the dLGN. H) Immunostaining of coronal dLGN sections from CyclinD2^WT^ and CyclinD2^cKO^ mice labeled with 488- and 594-CTB at P30. The entire dLGN, as well as the dLGN core receiving ipsilateral input, are outlined. Scale bar: 200μm. D: dorsal, V: ventral, M: medial, L: lateral. Axes indicate the orientation used for quantification along the DV and ML planes. I) Segregation plot: Percent of segregated inputs as a function of contralateral threshold. J) Area of the ipsilateral projection as percentage of total dLGN area. K) Extent of the ipsilateral projection as percentage of length along the DV and ML dLGN axes.

Because temporal control of cell birth is directly linked to the regulation of neurogenesis and cell fate specification, we tested whether CyclinD2 deficiency impedes cells from acquiring ipsilateral RCG identity through Zic2 expression. We noted a significant reduction in the number of Zic2 ^+^ RGCs in the CyclinD2^cKO^ ventrotemporal neural retina at E14.5 and E15.5, resembling the albino phenotype (Fig. 4E-F). We also probed for differences in contralateral RGCs between CyclinD2^WT^ and CyclinD2^cKO^ mice (Fig. S4B). We performed our measurements at E15.5, a time when Brn3a^+^ cells begin to be detected in the ventrotemporal neural retina among the Zic2^+^ population (Marcucci et al., 2018). CyclinD2 deletion from the CMZ did not lead to changes in the Brn3a^+^ population within this domain (Fig. S4C). These results are consistent with our finding that the ventral CMZ generates primarily Zic2^+^ RGCs (Fig. 1), and confirm the previously suggested (Marcucci et al., 2016) connection between defects in CyclinD2 expression and ipsilateral RGC neurogenesis in the albino visual system.

We next determined whether the decrease in Zic2^+^ RGCs in the CyclinD2^cKO^ retina is reflected in defects in afferent targeting of the dLGN, as described previously in albino mice. We used whole-eye CTB labeling (Fig. 4G) and traced RGC axons in target regions of the CyclinD2^WT^ and CyclinD2^cKO^ dLGN at P30. We found that both the ipsilateral and the contralateral retinogeniculate projections of CyclinD2^cKO^ mice appropriately innervate the core and the shell of the dLGN, respectively (Fig. 4H). However, the area receiving ipsilateral retinal input was markedly smaller in the CyclinD2^cKO^ dLGN compared with CyclinD2^WT^ (Fig. 4I-J). This effect extended along both the ML and the DV dLGN axes (Fig. 4K).

Thus, our results indicate that CyclinD2 is a crucial component of a CMZ-specific molecular machinery that coordinates cell cycle progression with ipsilateral RGC fate determination, and is necessary for proper retinogeniculate targeting.

### Depth perception requires CyclinD2 expression in the CMZ

We hypothesized that alterations in the proportions of ipsi- and contralaterally-projecting RGCs in albino and CyclinD2^cKO^ mice could lead to deficits in binocular vision. To test this, we adopted the binocularly-driven “ visual cliff” behavioral task, a well-established paradigm that measures depth perception in both humans and rodents (Lim et al., 2016). In this task, freely behaving mice can choose to step down from a stage to a platform that consists of a “ shallow side” directly displaying a black and white grid, and a “ deep side” displaying the same grid at 2.5ft depth from the platform’s surface and thus appearing as a “ drop-off” (Fig. 5A). We first compared the performance of pigmented and albino mice. Pigmented mice almost always stepped onto the shallow side, whereas albino mice failed to distinguish between the shallow and deep side of the platform (Fig. 5B, Figure S5A-B). Both pigmented and albino mice spent the same time making a decision to step on the platform (Fig. 5C). We next compared CyclinD2^WT^ and CyclinD2^cKO^ mice. Similar to albino mice, CyclinD2^cKO^ mice exhibited poor depth perception, indicated by the lower number of trials in which they stepped onto the shallow side and longer time to reach a decision, compared to CyclinD2^WT^ littermates that performed similarly to pigmented mice (Fig. 5D-E, Fig. S5C-D).

**Figure 5:**
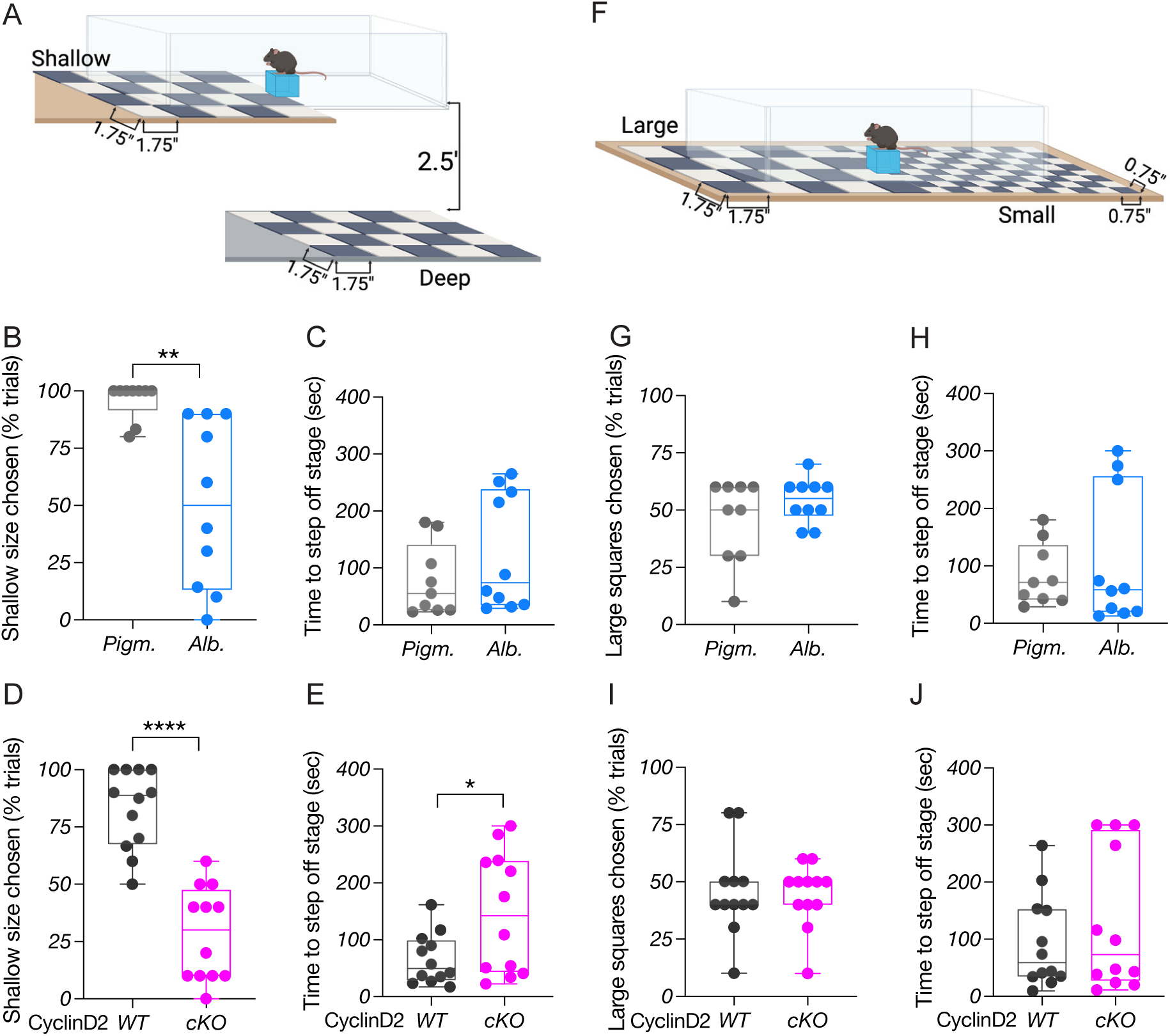
Depth perception requires CyclinD2 expression in the CMZ. A) Schema of the behavioral platform used for the binocularly-driven visual cliff assay. B) Quantification of depth perception, corresponding to the percentage of trials in which the shallow side was chosen, comparing pigmented and albino mice. C) Quantification of time (sec) needed to step off the stage to either side, comparing pigmented and albino mice. D) Quantification of depth perception, corresponding to the percentage of trials in which the shallow side was chosen, comparing CyclinD2^WT^ and CyclinD2^cKO^ mice. E) Quantification of time (sec) needed to step off the stage to either side, comparing CyclinD2^WT^ and CyclinD2^cKO^ mice. F) Schema of the behavioral platform used for the monocularly-driven relative size assay. G) Percentage of trials in which the large square-side was chosen, comparing pigmented and albino mice. H) Quantification of time (sec) needed to step off the stage to either side, comparing pigmented and albino mice. I) Percentage of trials in which large square-side was chosen, comparing CyclinD2^WT^ and CyclinD2^cKO^ mice. J) Quantification of time (sec) needed to step off the stage to either side, comparing CyclinD2^WT^ and CyclinD2^cKO^ mice.

To confirm that compromised depth perception in albino and CyclinD2^cKO^ mice resulted from ipsilateral RGC axon misrouting and binocular circuit abnormalities, we performed an additional visually-guided behavioral task that does not depend on binocular disparity, but instead relies on relative size, a monocular cue. For this task, both sides of the platform are at the same height, but its two sides display a low-versus a high-spatial frequency checkerboard pattern (Fig. 5F). Pigmented and albino mice behaved similarly in this monocularly-driven task, showing no preference to the low- or high-spatial frequency side (Fig. 5G). The time required to step from the stage to the platform was comparable between the two groups, and similar to their timing in the visual cliff task (Fig. 5H). The behavior of CyclinD2^WT^ and CyclinD2^cKO^ mice, regarding both choice and time-to-choice, was also similar during the relative size task (Fig. 5I-J). To further ensure that the behavior of CyclinD2^cKO^ mice was indicative of reduced depth perception, and not due to other visual deficits that could prevent them from discriminating the black-and-white grids clearly, we tested CyclinD2^WT^ and CyclinD2^cKO^ mice for contrast sensitivity and visual acuity using the Optodrum (Fig. S5G). Both groups, were able to track the stimulus until its contrast dropped to 4.92% (Fig. S5H), and had average visual acuity 0.45 cycles/degree (Fig. S5I).

Together, our results show that perturbations in the ratio of ipsi- to contralateral RGC projection leads to poor depth perception both in albino and CyclinD2^cKO^ pigmented mice, indicated by their inferior performance in the visual cliff task but not in the relative size task. Thus, CyclinD2 expression within the CMZ, and its involvement in regulating ipsi-/contralateral RGC neurogenesis, is critical for establishing the circuit for stereopsis.

### CyclinD2 upregulation restores binocular circuit formation and function in albino mice

Given the instrumental role of CyclinD2 in the proper formation of the binocular circuit, we asked whether we could rescue the reduced ipsilateral RGC population and impaired visually-guided behavior of albino mice by normalizing CyclinD2 expression in the albino CMZ. To upregulate CyclinD2 we injected into pregnant dams the L-type calcium channel agonist BayK-8644, which can increase CyclinD2 protein levels in the pancreas (Salpeter et al., 2011), and used sham-injected mice as controls. The use of BayK was appropriate because the *Cacna1d* gene and its encoding protein Cav1.3, which forms the pore subunit of the calcium channel, is expressed within the neurogenic (Msx1^+^) domain of both the pigmented and albino CMZ (Fig. S6A-B). L-type calcium channel activation can induce *Fos, Fosb*, and *Jun* (Cruzalegui et al., 1999; Leclerc et al., 1999). Our scRNA-Seq DE analysis revealed a significant downregulation of these three genes in the albino CMZ (Fig. S6C-E). Importantly, *Fos, Fosb*, and *Jun* can act as transcriptional regulators of CyclinD proteins, suggested by studies in other cell types (Brown et al., 1998; Turchi et al., 2000).

Administration of BayK at E13.5 and E14 elevated the expression of CyclinD2 in the albino ventral CMZ at E15.5, bringing it to pigmented-like levels (Fig. 6A-C). In addition to CyclinD2, BayK led to a significant increase of Zic2^+^ RGCs in the ventrotemporal retina, compared to sham-treated albino mice (Fig. 6D-E). However, BayK had no effect on the Zic2^+^ RGC population of CyclinD2^cKO^ mice, which remained diminished compared to CyclinD2^WT^ (Fig. S7A-B) suggesting that the BayK-induced Zic2^+^ RGC augmentation in the albino retina requires CyclinD2.

**Figure 6:**
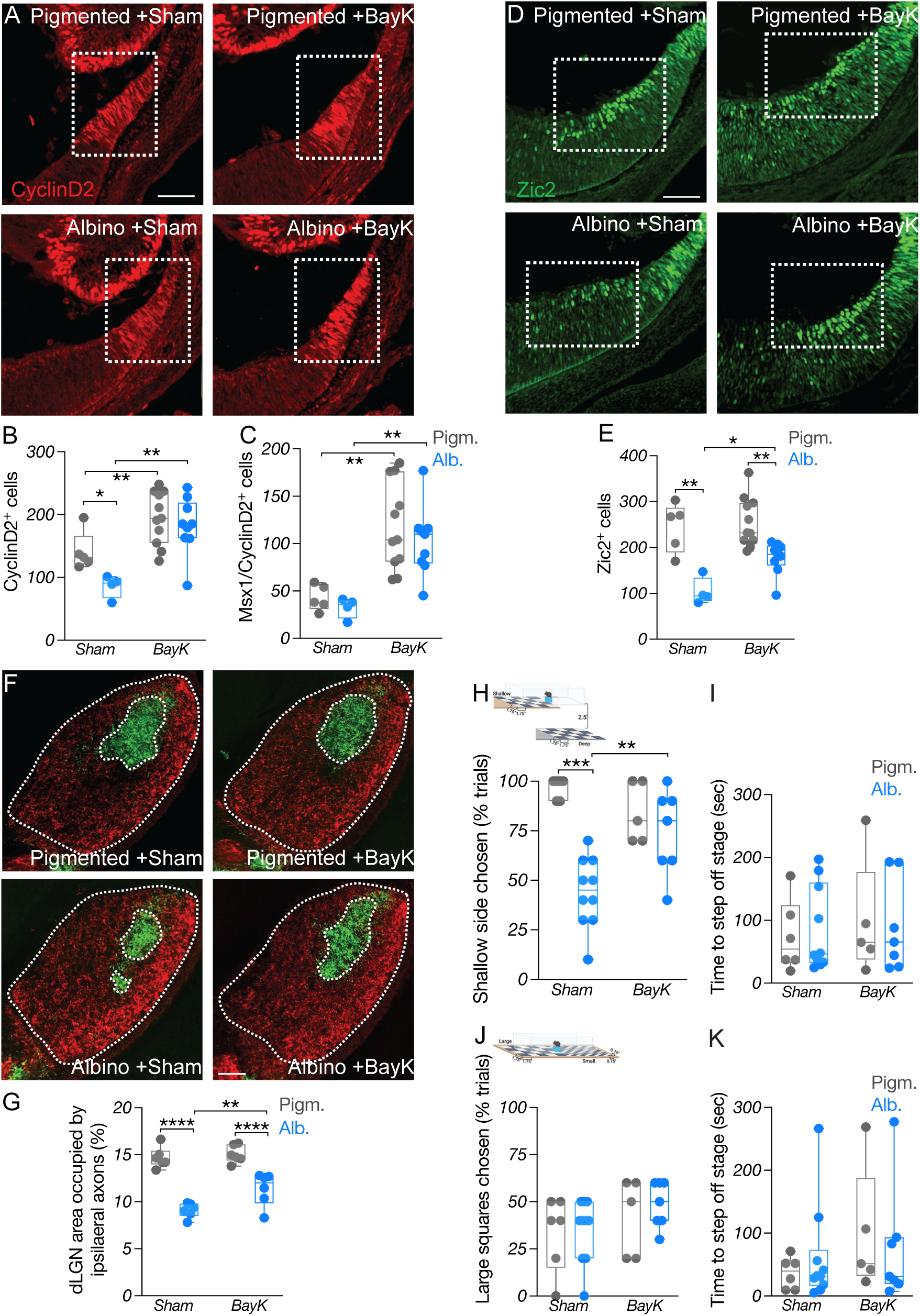
CyclinD2 upregulation restores binocular circuit formation and function in albino mice. A) Immunostaining of CyclinD2 in the ventrotemporal CMZ of pigmented and albino Sham vs BayK treated mice at E15.5. Scale bar: 100μm. B-C) Quantification of CyclinD2^+^ (B) and CyclinD2^+^/Msx1^+^ (C) cells at E15.5. D) Immunostaining of Zic2 in the ventrotemporal retina of pigmented and albino Sham vs BayK treated mice at E15.5. Scale bar: 100μm. E) Quantification of Zic2^+^ cells at E15.5. F) Immunostaining of coronal dLGN sections from pigmented and albino Sham vs BayK treated mice labeled with 488- and 647-CTB at P60. The entire dLGN, as well as the dLGN core receiving ipsilateral input, are outlined. Scale bar: 100μm. G) Area of the ipsilateral projection as percentage of total dLGN area. H) Depth perception, corresponding to the percentage of trials in which the shallow side was chosen during the visual cliff task, comparing pigmented and albino Sham vs BayK treated mice. I) Quantification of time (sec) needed to step off the stage to either side during the visual cliff task. J) Percentage of trials in which large square-side was chosen during the relative size task, comparing pigmented and albino Sham vs BayK treated mice. K) Quantification of time (sec) needed to step off the stage to either side during the relative size task.

We then used whole-eye CTB labeling to assess whether the ipsilateral innervation of the albino dLGN was also ameliorated upon BayK treatment. As expected, the dLGN core area receiving ipsilateral retinal input was notably smaller in sham-treated albino mice compared to pigmented mice. This defect was partially reversed in BayK-treated albino mice, in which the ipsilateral retinorecipient region of the dLGN was significantly increased compared to the sham albino group (Fig. 6F-G).

To test whether the restoration of the ipsilateral retinogeniculate projection ultimately leads to improved depth perception, we subjected sham- and BayK-treated pigmented and albino mice to the visual cliff and relative size behavioral tasks. Albino mice that received a sham treatment during development exhibited poor depth perception compared to pigmented littermates, as expected. Strikingly, albino mice that received BayK chose mostly the shallow side of the visual cliff, performing similarly to pigmented mice (Fig. 6H-I and Fig. S7C). In contrast to their behavior during the visual cliff, which was guided by binocular cues, all four groups of mice displayed similar behavior during the monocularly-driven relative size task (Fig. 6J-K and Fig. S7D).

Lastly, we tested whether the BayK-induced improvement in depth perception could be observed in albino mice of another genetic background. We thus treated Swiss albino mice with sham or BayK during development and tested their adult visually-guided behavior. Similar to Tyr^c2j^ albino mice, Swiss albinos that received BayK exhibited considerably improved behavior in the visual cliff test compared to those that received sham (Fig. S7E-F), while both groups performed similarly in the relative size task (Fig. S7G-H).

In summary, our results revealed a CMZ-specific mechanism that controls RGC neurogenesis and ipsilateral fate in the ventrotemporal retina, where ipsilateral RGCs come to reside, and showed that pharmacological stimulation of this mechanism can restore the binocular circuit and depth perception in albinism (Fig. 7).

**Figure 7:**
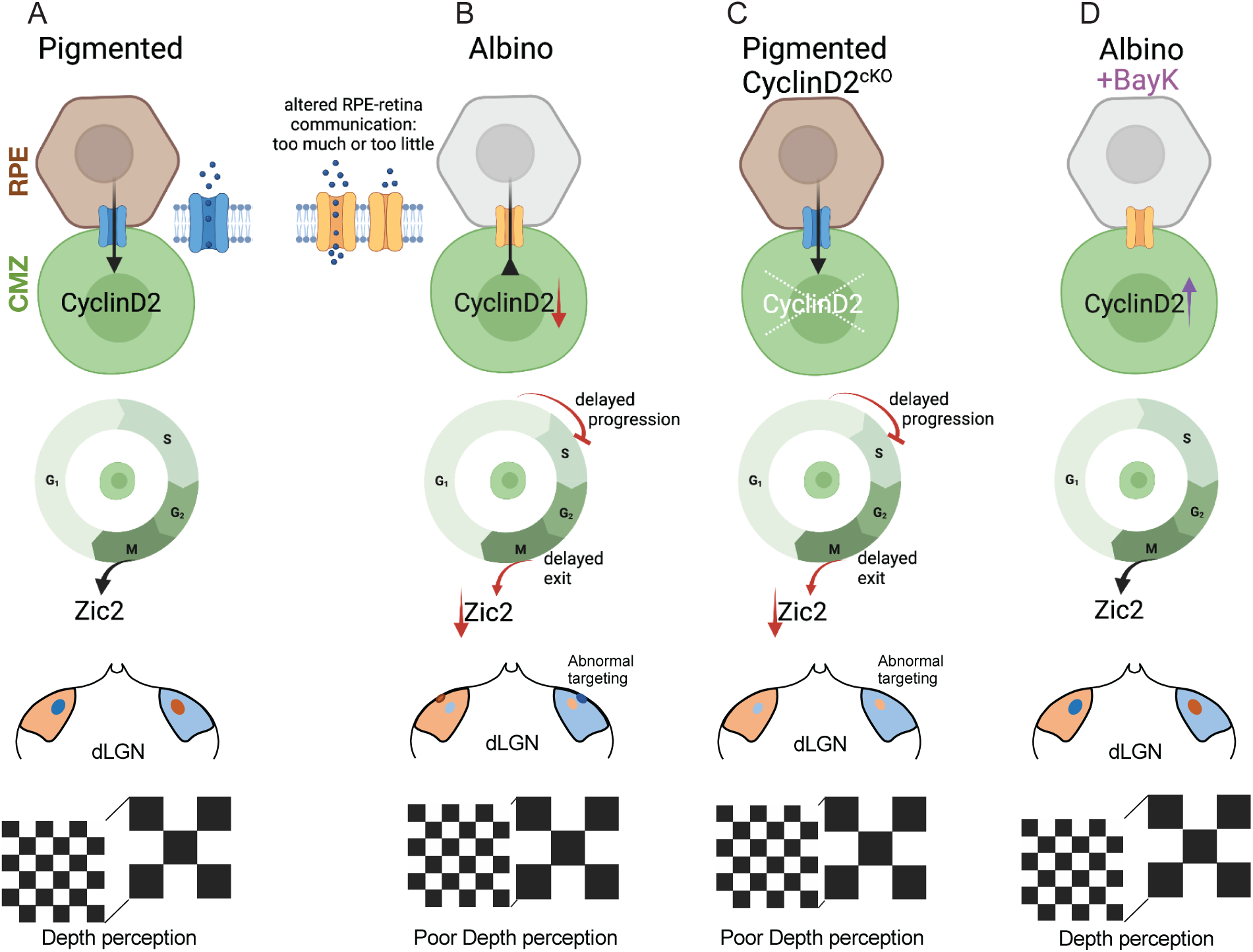
Summary. A) The link between CyclinD2 and cell cycle, ipsilateral RGC hallmark gene (Zic2), retinogeniculate projection, and binocular vision. B) In the albino, CyclinD2 reduction leads to delay in cell cycle progression from G1 to S phase, reduced numbers of Zic2^+^ RGCs, reduced ipsilateral retinogeniculate projection, and poor depth perception. C) This phenotype is recapitulated when CyclinD2 is deleted from the pigmented peripheral retina. D) The binocular circuit-related deficits in the albino retina can be rescued by the delivery of the L-type calcium channel agonist BayK-8644.

## Discussion

In this study, we focus on the mouse CMZ and discern a cellular mechanism regulating the ipsi-/contralateral output of RGC neurogenesis from this zone, that is defective in the albino visual system, thereby leading to poor binocular vision.

### The ciliary margin zone as a source of retinal neurons

During retinal development, most neurons are generated following a center-to-periphery gradient. In lower species such as fish and amphibians, retinal neurons continue to arise over the lifespan from the CMZ, a highly proliferative zone in the retinal periphery (Centanin et al., 2011; Fischer and Reh, 2000; Moshiri and Reh, 2004; Perron and Harris, 2000; Wan et al., 2016; Wetts et al., 1989). In mammals, the CMZ has mostly been studied in the context of iris and ciliary body formation, whereas its neurogenic potential has been debated (Fernandez-Nogales et al., 2019). It is commonly believed that the adult postnatal CMZ is unable to generate neurons, thereby precluding the mammalian retina from having a regenerative capacity (Kubota et al., 2002). In contrast to these reports, a proliferating population of CMZ stem cells has been described both *in vitro* (Tropepe et al., 2000) and in adult mice carrying a Shh perturbation (Moshiri and Reh, 2004). Similar to CMZ neural precursors seen in lower adult vertebrates, these cells have the potential to differentiate into retinal cells (Tropepe et al., 2000). Supporting these studies, cells of the embryonic CMZ have been shown to migrate laterally to the adjacent neural retina, where differentiated RGCs reside (Marcucci et al., 2016). Additionally, cells that are born in the embryonic mouse CMZ and express the transcription factor Msx1 have been detected in the postnatal neural retina, where they differentiate into various neuronal subtypes, including RGCs (Belanger et al., 2017).

Our research builds upon these findings and enlarges our view of the neurogenic role of CMZ, by revealing the identity of CMZ-derived RGCs. Our fate mapping experiments, distinguishing between ventral and dorsal segment, show that the ventral CMZ is competent to generate ipsilateral RGCs, supplementing the ventrotemporally-confined pool of ipsilateral neurons born via center-to-periphery neurogenesis. It remains to be determined whether the CMZ produces contralateral RGCs at later time-points not tested here (i.e., E17.5 until birth) and whether, in contrast to the ventral CMZ, the dorsal CMZ produces contralateral RGCs. Nevertheless, our results raise the question of whether progenitors within the CMZ are intrinsically distinct from retinal progenitors, in that they have lost their multipotency and are committed to the ipsilateral fate while still in the cell cycle. We also show that the albino retina bears CMZ-specific disruptions that prevent the generation of ipsilateral RGCs, strongly suggesting that the anomalies of the binocular circuit in albinism are rooted in the CMZ niche. This finding provides further evidence that RGC neurogenesis from the CMZ is indispensable to the proper formation of the binocular circuit.

### Transcriptional profiles of ipsi- and contralateral RGCs

Single-cell profiling studies have illuminated the transcriptional programs conveying diverse neuronal identities in the human and mouse retina (Clark et al., 2019; Lo Giudice et al., 2019; Lu et al., 2020; Shekhar et al., 2021; Sridhar et al., 2020; Trakhtenberg et al., 2018). Although the involvement of CMZ-specific genes in retinal neurogenesis has been overlooked, despite a recent CMZ transcriptomic analysis (Balasubramanian et al., 2021), two independent observations support the role of CMZ as a neuronal source. First, a small group of cells arising from postmitotic neuroblasts in the ciliary margin have been found positive for RGC markers (Lo Giudice et al., 2019). Second, an early neuroepithelial progenitor cell cluster has been shown to express the CMZ genes *Ccnd2* and *Msx1* (Clark et al., 2019). These reports support our findings implicating CyclinD2 in neurogenesis from Msx1^+^ progenitors of the CMZ.

Much has also been learned about regulatory networks driving the acquisition of ipsi- and contralateral RGC properties. Ipsilateral identity in RGCs of the ventrotemporal retinal crescent is specified by the expression of *Zic2* (Herrera et al., 2003). *Igfbp5, Tc1, Sst, Gal, Crabp1, Pcsk2, Lgi2*, and *Zic5* expression coincides with *Zic2*, and thus these genes were recently linked to ipsilateral identity. *SoxC*s, *Isl2, Fgf12, Pou3f1, Igf1, Nrp1, Lmo2, Pcsk1n*, and *Syt4* are all enriched in contralateral RGCs (Lo Giudice et al., 2019; Shekhar et al., 2021; Wang et al., 2016). Our study provides insights on how the expression of these genes differs in the albino retina, in conjunction with the reduced ipsilateral RGC population. In addition to *Zic2, Igfbp5* and *Pcsk2* were downregulated in neurogenic and RGC albino clusters, respectively. In contrast, the contralateral genes *Pou3f1, Nrp1*, and *Fgf12* were upregulated in albino postmitotic RGCs. It appears that *Zic2* and *Igfbp5* expression in the pigmented peripheral retina begins in unspecified cells and gradually wanes during RGC maturation. Whether *Zic2* drives the expression of *Igfbp5* or vice versa has not been determined. Nevertheless, at the late neurogenic stages, these genes may repress contralateral determinants to promote ipsilateral fate. This repression is not achieved in the albino, due to low *Zic2* and *Igfbp5* levels, potentially allowing the acquisition of contralateral identity in the ventrotemporal crescent.

### Temporal regulation of neurogenesis and ipsilateral cell fate

The specification of the different retinal cell types and subtypes is directly related to the timing of cell cycle exit (Cepko et al., 1996; Marcucci et al., 2018; Osterhout et al., 2014). Thus, the generation of different RGC populations is chronologically distinct: ipsilateral (Zic2^+^) RGCs peak in the ventrotemporal retina at E14.5, whereas contralateral RGCs are born in the rest of the retina by E16.5 and in the ventrotemporal retina from E15.5 onward (Herrera et al., 2003; Marcucci et al., 2018). In the albino visual system, RGC neurogenesis at E14.5 is reduced in the ventrotemporal segment. Although this reduction is compensated by RGC overproduction at later stages, suggesting a shift in the timing of neurogenesis, the Zic2-expresing population is inevitably smaller (Bhansali et al., 2014).

Tightly regulated birth timing, and therefore cell fate specification, relies on the expression of specific cell cycle regulatory molecules. Together with our birthdating experiments, our scRNA-Seq analysis in the pigmented and albino peripheral retina, revealed that the developmental trajectories of CMZ-derived RGCs are temporally controlled by CyclinD2. This is in agreement with previous studies reporting reduced CyclinD2 expression in the albino retina (Marcucci et al., 2016) and with experiments in global CyclinD2-knockout mice implicating CyclinD2 deficits in reduced birth of ipsilateral RGCs (Marcucci et al., 2018). Our results obtained both from albino and conditional CyclinD2-deficient pigmented mice go further than previous studies and show that CyclinD2 downregulation leads to delayed G1-to-S phase transition and mitotic exit of albino CMZ progenitors. This perturbation in the temporal control of the cell cycle by CyclinD2 coincides with a time window that is crucial for ipsilateral RGC neurogenesis (E13.5-14.5), and therefore results in reduced Zic2^+^ RGC production from the CMZ. Ultimately, the architecture of the binocular circuit is altered due to a diminished ipsilateral retinogeniculate projection, leading to compromised depth perception.

Although conditional CyclinD2-deficient pigmented mice largely recapitulate the albino phenotype in several aspects of cell cycle and mitotic exit, Zic2^+^ RGC neurogenesis, and binocular circuit architecture and function, differences between the two do exist. Namely, the reduction in the Zic2^+^ RGC population is less severe in the CyclinD2^cKO^ compared to the albino, despite the complete absence of CyclinD2 from the CMZ in the former. Similarly, the thalamic area receiving ipsilateral retinal input is less impacted in CyclinD2^cKO^ compared to albino mice. These observations favor the hypothesis that the mouse retina is governed by two distinct routes of ipsilateral RGC neurogenesis, one that follows the well-established center-to-periphery direction, and one that originates in the CMZ and is controlled by CyclinD2. However, perturbed melanin biogenesis in the albino retinal pigment epithelium (RPE) can influence these routes of neurogenesis in ways that extend beyond the loss of CyclinD2 in the conditional knockout.

### Depth perception in albino mice

The primate visual system relies on binocular disparity-based stereopsis to calculate depth (Cumming and DeAngelis, 2001; Parker, 2007), thereby enabling a three-dimensional view of the environment. Binocular disparity-tuned neurons have also been characterized in the mouse visual cortex, enabling the discrimination of stereoscopic depth (Samonds et al., 2019), which in turn guides behavior both in natural scenes, i.e., during prey approach and contact (Johnson et al., 2021), and in non-natural settings, i.e., during a pole descent cliff assay (Boone et al., 2021). The cliff response is one of the few innate visual behaviors in mice (Fox, 1965). In our study, we use the visual cliff behavioral task to measure depth perception, and to determine whether the functional integrity of the albino retinogeniculate circuit is compromised due to aberrations in its architecture. Similar to observations in albino rats (Walk and Gibson, 1961), we find that both albino and pigmented CyclinD2-deficient mice perform poorly in the visual cliff paradigm. Our result opposes an older study showing that depth perception differences in albino mice of different strains are not detected by the visual cliff assay (Fox, 1965). We argue that strain-related differences could underlie this discrepancy. For example, light or contrast sensitivity, visual acuity, or other important factors may vary across albino strains. Such differences, if they exist, would influence the view of the behavioral stimulus as a whole, not just the aspect of depth. To fully address the issue of depth perception albinism and its impact on the interactions with the surrounding environment, it would be of interest to characterize the behavior of these mice during other visually-guided tasks, such as prey hunting (Johnson et al., 2021). It would also be relevant to address whether and how neuronal ensembles responsible for depth perception are differentially activated in the visual cortex of mice with misrouted retinogeniculate pathway due to CyclinD2-deficiency.

### The role of the retinal pigment epithelium in retinal development

RPE cells physically and functionally interact with each other and with the adjacent neural retina to ensure its proper development and structural organization (Amram et al., 2017; Martinez-Morales et al., 2004). Delays in center-to-periphery neuronal differentiation (Bhansali et al., 2014), aberrant patterns of retinal cell division, differences in mitotic spindle orientation, and failure to transition from symmetric/proliferative to asymmetric/neurogenic cell division modes (Cayouette et al., 2001; Ilia and Jeffery, 1996; Tibber et al., 2006), have been associated with absence of RPE and/or pigmentation. Albino RPE cells themselves exhibit atypical morphological and biological features, including the gap junctions that couple RPE cells with the adjacent apical surface of the neural retina where neuroblasts divide (Iwai-Takekoshi et al., 2016). To this day, the link between perturbed melanogenesis in the RPE and aberrations in the circuitry from the eye to the brain seen in albinism remains unresolved. One explanation suggested by the present data is that the RPE provides signals to the neural retina that regulate the tempo of the cell cycle during neurogenesis and fate specification of RGCs. In albinism, RPE-originating signals may be either insufficient or excessive, due to metabolic and cellular aberrations caused by the lack of pigment (Fig. 7). A possible cause of disrupted RPE-neural retina signaling in the embryonic albino eye is the downregulation of L-Dopa (Roffler-Tarlov et al., 2013), a tyrosine derivative that has been proposed to influence the pace of the cell cycle and the plane of cell division (Kralj-Hans et al., 2006; Lavado et al., 2006) and has been identified as the ligand for GPCRs on the melanosome surface (Lopez et al., 2008).

Our current work points to calcium as another potential signal that influences the output of CMZ neurogenesis. Melanin in the RPE has calcium buffering capacity and has been associated with the tight regulation of calcium levels available to the neural retina (Ambrosio et al., 2016; Bellono and Oancea, 2014; Bush and Simon, 2007; Drager, 1985; Salceda and Sanchez-Chavez, 2000). In turn, calcium transients have been shown to regulate retinal development by inducing progenitor proliferation (Pearson et al., 2002). Here, stimulation of L-type calcium channels with BayK-8644 to pregnant dams, led to markedly increased CyclinD2 expression, a larger Zic2^+^ RGC population in the albino retina, and ultimately improved function of the binocular circuit. How calcium triggers CyclinD2 expression calls for further investigation. According to our scRNA-Seq data, *Fos, Fosb*, and *Jun*, which have been described as upstream regulators of the expression of CyclinD proteins (Brown et al., 1998; Turchi et al., 2000) are downregulated in the albino CMZ. BayK-mediated calcium influx may induce the transcription of *Fos, Fosb*, and *Jun* (Cruzalegui et al., 1999; Leclerc et al., 1999) in cells of the albino CMZ, where Cav1.3 channels are present, thereby augmenting the levels of CyclinD2.

Collectively, our work highlights CyclinD2 as a key regulator of the neurogenic output of the CMZ by controlling aspects of the cell cycle, and thus the timely expression of transcription factors specific to ipsilateral RCG fate. In addition, our results establish a critical link between calcium channel function and RGC fate acquisition, providing an inroad to unravel the relationship between perturbed melanogenesis in the RPE and aberrations in the circuitry from the eye to the brain seen in albinism.

### Outlook

Optic nerve degeneration leads to RGC death, while the lack of spontaneous regenerative response of the retina results in permanent damage of central visual pathways between the retina, LGN and visual cortex. In the past decade, remarkable advances in post-injury RGC survival and axon regeneration, as well as in stem cell-based replacement therapies, offer great promise for the repair and recovery of the damaged visual system. Nonetheless, integration of RGCs after their transplantation to the host tissue, as well as directed axon growth through and past the optic chiasm, pose significant challenges. The CMZ has stem-cell like capacity under the control of Fgf signaling (Balasubramanian et al., 2021). The CMZ may therefore have high therapeutic value as a niche of endogenous retinal progenitor cells, which can be expanded *in vitro* and differentiated into RGCs. Thus, future studies on the neurogenic properties of the developing CMZ, the temporal parameters of progenitor-to-RGC progression, and the directives for specification of RGCs into ipsi- and contralateral subtypes, can be leveraged to develop stem cell-based approaches for successful restoration of the visual pathway in injuries or sight-threatening conditions.

## Acknowledgements

We thank Ira Schieren in the Flow Cytometry Platform, Susan Morton in the Antibody Platform at the Zuckerman Institute, and the Sulzberger Genome Center at Columbia University. Imaging was performed with support from the Zuckerman Institute’s Cellular Imaging Platform. This work also used the Genomics and High Throughput Screening Shared Resource. We also thank Xin Zhang (Dept. of Ophthalmology, Columbia University) for providing the Msx1 Cre and aCre mice, Stephen Brown (University of Vermont) for providing the Zic2 antibody, Sania Khalid for technical help, and Jane Dodd and Xin Zhang for reading the manuscript. This work was funded by NIH R01 EY015290, R01 EY12736, António Champalimaud Vision Award, Simons Foundation Senior Fellow Award, and Vision of Children (CAM), NIH R01 EY032062 and R01 EY032507 (SWMJ), Precision Medicine Initiative at Columbia University (SWMJ), R01 NS105477 (MER), Vision Core Grant P30 EY019007 (M. Goldberg), NIH/NCI Cancer Center Support Grant P30CA013696 and National Center for Advancing Translational Sciences, National Institutes of Health, through Grant Number UL1TR001873 (Sulzberger Genome Center). Financial support from Fight for Sight and a Georgakopoulos Family Fellowship Award to NS is gratefully acknowledged. RB is supported by the BrightFocus Foundation Grant G2021007S.

## Methods

### Animals

Mice were maintained under standard conditions approved by the Institutional Animal Care and Use Committee (IACUC) of Columbia University. Animals were housed in a specific pathogen– free animal house under a 12/12-hour light/dark cycle and were provided food and water. Msx1-Cre^ERT2^ mice in C57Bl/6J background and αCre mice on a mixed genetic background were a gift from Dr. Xin Zhang, Columbia University. αCre mice were backcrossed for several generations with wild type C57Bl/6J mice (000664, Jackson Laboratories). CyclinD2^flox^ mice on a C57Bl/6J background were a gift from Dr. Elizabeth Ross, Weill Cornell. tdTomato Ai14 (007914), Tyr^c2j^ (000058), and Swiss/ICR albino (034608) mice were purchased from Jackson Laboratories. Tyr^c2j^ mice were bred with wild type C57Bl/6J mice to obtain pigmented Tyr^WT/c2j^ controls. Animals of both sexes were used in our experiments.

### Anesthesia

Ketamine and Xylazine were diluted in saline, and administered at 100mg/kg and 10mg/kg, respectively. Toe pinch was used to confirm full anesthesia.

### Single cell dissociation and sorting

We used αCre; tdTomato^+^ mice, in which the retinal periphery including the CMZ is fluorescently labeled upon activation of the retina-specific regulatory element of Pax6 α-enhancer (Marquardt et al., 2001), to enrich CMZ cells in our samples. E13.5 retinae with attached RPE from 4 pigmented and 5 albino αCre; tdTomato^+^ littermate embryos each were harvested in ice cold Dulbecco’s modified Eagle’s medium (DMEM) after removal of the lens. Dissociation was performed in activated papain solution including Deoxyribonuclease I, using a papain dissociation system (Worthington Biochemical) for 10 min at 37°C and stopped using DMEM + 10% fetal bovine serum (FBS). Cells were gently triturated using a 22G needle, centrifuged at 300g for 5min, and washed with DMEM + 10% FBS. After passing through a 70μm filter and washing again in DMEM + 10% FBS, the cell suspension was stained with DAPI (0.2μg/ml) to label dead cells. Flow cytometry was performed at the core facility of Zuckerman Institute, Columbia University. Single cells were collected with an Astrios cell sorter and gated for live cells and TdTomato^+^ cells using FlowJo software. Approximately 25,000 pigmented and 15,000 albino cells were collected into 1.5-ml tubes precoated with FBS and DMEM + 10% FBS for submission for single-cell sequencing.

### scRNA-Seq

Single-cell sequencing was performed at the single-cell sequencing core in Columbia Genome Center. Flow-sorted single cells were loaded into chromium microfluidic chips with v3 chemistry and barcoded with a 10× chromium controller (10× Genomics). RNA from the barcoded cells was reverse-transcribed, and sequencing libraries were constructed with a Chromium Single Cell v3 reagent kit (10× Genomics). Sequencing was performed on NovaSeq 6000 (Illumina).

Raw reads mapped to the mm10 reference genome by 10× Genomics Cell Ranger pipeline (v2.1.1) using default parameters were used for all downstream analyses using Seurat v3 (Stuart et al., 2019), Velocyto and scVelo. Briefly, the dataset was filtered to contain cells with at least 200 expressed genes and genes with expression in more than three cells. Cells were also filtered for mitochondrial gene expression (<30%). The dataset was log-normalized and scaled. Unsupervised clustering was performed initially, followed by manual annotation of Seurat clusters. An integration analysis was performed to compare and analyze the control and mutant gene expression matrices, which included normalization for cell numbers. The unsupervised clustering with a resolution parameter of 0.5 for both control and mutant cells was represented on a UMAP space, and cluster identity was assigned. Expression of various known genes was used to determine cluster identities. Doublets identified based on biological incompatibility were manually excluded from the dataset. Extra-retinal contaminant cells were manually removed from the dataset. The cell cycle was assessed using the vignette from Seurat. For RNA velocity analysis, loom files were constructed by extracting the reads from the 10× single-cell dataset. After gene filtering, spliced and unspliced counts were normalized based on total counts per cell. Velocity plots are represented as the transcriptomic integration of control and mutant datasets to represent similarities and differences in their velocities, respectively. Transcriptional dynamics were determined by ratio of spliced to unspliced counts fitted to a generalized linear regression model (Bergen et al., 2020; La Manno et al., 2018).

### Intraperitoneal injections into pregnant dams

4-Hydroxytamoxifen (4-OHT) (Tocris Bioscience) was dissolved in corn oil (Millipore-Sigma) at final concentration 5mg/ml and was administered intraperitoneally at 25μg/g body weight. BrdU (Thermo Fisher Scientific) was dissolved in 10mM Tris-HCl pH7.6 (Millipore-Sigma) at stock concentration 40mg/ml. Prior to injection, stock was further diluted in PBS and administered at 100μg/g body weight. EdU (Thermo Fisher Scientific) was dissolved in PBS at stock concentration 2.5mg/ml. Prior to injection, stock was further diluted in PBS and administered at 10μg/g body weight. BayK-8644 was dissolved in 1ml DMSO (Millipore-Sigma) at stock for concentration 5mg/ml. Prior to injection, stock was further diluted 1:10 in saline and administered at 2 mg/kg body weight. DMSO diluted 1:10 in saline was used as sham, at the same injection volume as BayK-8644.

### Fate mapping

Three doses of 4-OHT were administered intraperitoneally to Msx1-Cre^ERT2^ mice; tdTomato^+^ pregnant dams at E13.5, E14.5, and E15.5. EdU was also administered at E15.5. Embryos were collected at E17.5 and processed for immunohistochemistry.

### Dual pulse birthdating

At the day of the experiment (E13.5 or E14.5), BrdU was injected intraperitoneally into the pregnant dam. 2 hours later EdU was administered in the same route. Embryos were collected 30min later and processed for immunohistochemistry. Cell cycle duration (hours) was calculated using the formula: Tc = 2*(Ki67^+^cells / BrdU only^+^ cells)

### Anterograde labeling of retinogeniculate projections

Cholera toxin subunit B conjugated to Alexa Fluor 488 or 594 (Thermo Fisher Scientific) was dissolved in PBS with 0.1% DMSO at stock concentration 1mg/ml. Mice were fully anesthetized and 1-2μl of CTB were injected intravitreally with a glass micropipette, until the whole eye was filled. Mouse brains were processed 48 hours later.

### Tissue processing for cryosections

Pregnant dams were anesthetized and embryos were collected in ice cold PBS. E13.5-E15.5 embryo heads were fixed in 4% paraformaldehyde (PFA) (Electron Microscopy Sciences) in PBS for 2 hours at 4°C. E17.5 embryos were transcardially perfused with cold 4% PFA and heads were post-fixed in 4% PFA overnight at 4°C. PFA was rinsed with PBS, and heads were incubated in 10% sucrose in PBS overnight at 4°C, followed by 20% sucrose in PBS overnight at 4°C, and 30% sucrose in PBS overnight at 4°C. Heads were cryomolded in O.C.T. Compound (Thermo Fisher Scientific) in dry ice and stored at -80°C. Coronal sections were obtained at 12μm thickness on Superfrost microscope slides (Thermo Fisher Scientific).

### Tissue processing for brain floating sections

Adult mice were transcardially perfused with 10ml cold PBS followed by 30ml cold 4% PFA, and brains were dissected and post-fixed in 4% PFA overnight at 4°C. PFA was rinsed with PBS, and brains were molded in 3% agarose in PBS. Vibratome sections were obtained at 150μm thickness in ice cold PBS and mounted on Superfrost microscope slides.

### Immunohistochemistry and Imaging

Frozen sections were rinsed with PBS. Antigen retrieval was performed in an 85°C water bath for 15 minutes using pre-heated sodium citrate+0.05% Tween20, pH6. Slides remained incubated in the sodium citrate solution at room temperature for 20min, then were washed three times with PBS and incubated with blocking solution of 10% normal donkey serum (Jackson immunoResearch) in PBS with 0.2% Tween20, for 1 hour at room temperature inside a humid chamber protected from light. After blocking, primary antibody solution (1% NDS in PBS-T) was applied overnight at 4°C. Primary antibodies used were: rabbit anti-Zic2 (1:10,000; gift from Dr. Steve Brown, University of Vermont), mouse anti-Islet1/2 (1:100; gift from Susan Morton, Columbia University), guinea pig anti-RFP (1:30,000; gift from Susan Morton, Columbia University), mouse anti-Brn3a (1:50; EMD Millipore), rat anti-CyclinD2 (1:50; Santa Cruz Biotechnology), rat anti-Ki67 (1:100; Thermo Fisher Scientific), mouse anti-BrdU (1:200; Thermo Fisher Scientific), rabbit anti-phosphoHistone3 (1/300; Thermo Fisher Scientific), goat anti-Msx1 (1:100, R&D systems), mouse anti-Cav1.3 (1:100, GeneTex), mouse anti-Cre recombinase (1:100, Millipore-Sigma). Primary antibody solution was washed three times with PBS-T and secondary antibody solution (1:500 in 1% NDS in PBS-T) was applied for 2 hours at room temperature, protected from light. Secondary antibodies used were: donkey anti-rat AlexaFluor594, donkey anti-mouse AlexaFluor647, donkey anti-mouse AlexaFluor405, donkey anti-mouse AlexaFluor488, donkey anti-rabbit AlexaFluor488, donkey anti-rabbit AlexaFluor647, donkey anti-guinea pig AlexaFluor594, donkey anti-goat AlexaFluor647, donkey anti-goat 488. All secondary antibodies were purchased from Jackson ImmunoResearch. Secondary antibodies were washed three times with PBS. For EdU visualization, Click-iT reaction was performed after secondary antibody incubation, for 30min at room temperature according to kit instructions (Thermo Fisher Scientific). Slides were incubated with DAPI (Fisher Scientific) 1:1,000 in PBS for 10min at room temperature, and mounted using Fluoro-Gel Medium (Electron Microscopy Sciences). Fluorescent imaging was performed on a W1-Yokogawa spinning disk confocal microscope. Images for Z-stacks were obtained at 0.5μm step size.

### Cell number quantification

The CMZ in each section was defined as the region extending 150μm from the distal-most tip of the retina. Cells were counted manually using the cell counter plugin in Fiji software. Individual cells were distinguished by DAPI staining. In the entire series of sections, the first one that included the optic nerve head opening was assigned as “ 0μm distance from the optic nerve head” . Because sections were 12μm thick, every section along the rostro-caudal axis was assigned as “ n+12μm distance from the optic nerve head”, with n being the distance of the previous section. The counts obtained from all sections of the temporal retina were summed. For E13.5 and E14.5 eyes, 6 serial sections were analyzed, starting from the one at 0μm distance from the optic nerve head. For E15.5 eyes, 9 serial sections were analyzed, starting from the one at 0μm distance from the optic nerve head. For E17.5 eyes, 32 serial sections were analyzed, starting from the one at 0μm distance from the optic nerve head. Left and right eyes were aligned based on distance from the optic nerve head and their counts were averaged.

### Analysis of ipsilateral input to the dLGN

Quantification of eye-specific axon terminals was performed on 10x fluorescent images from three consecutive coronal 100μm sections through the region of the dLGN containing the greatest extent of the ipsilateral projection, as described previously (Rebsam et al., 2012). The extent of eye-specific projections within the dLGN along the dorsoventral (DV) and mediolateral (ML) axes of the dLGN was measured by tracing a line along each axis of the dLGN. The extent of the ipsilateral projection was measured by tracing a line between the two points delineating the maximal extent of the ipsilateral signal along the chosen axis. The length of the ipsilateral projection was divided by the length of the dLGN for each axis. To calculate the area of the dLGN receiving ipsilateral input we traced the boundary of the dLGN was outlined, excluding the intrageniculate leaflet, the ventral lateral geniculate nucleus, and the optic tract, as well as the boundary of the central ipsilateral patch, and identified the pixels included in each traced region. The pixels included in the traced ipsilateral region were divided by the pixels included in the traced dLGN region. We conducted an additional eye-specific segregation analysis, using MetaMorph software (Molecular Devices). We used variable thresholds for the contralateral projection and a fixed threshold for the ipsilateral projection. The boundary of the dLGN was outlined, as described above. The intensity threshold for the ipsilateral projection was chosen when the signal-to-background ratio was at least 1.2. The overlap between ipsilateral and contralateral projections (pixel overlap) was measured at every 10th threshold value for the contralateral image to obtain the function of overlap between ipsilateral and contralateral projections. The proportion of dLGN occupied exclusively by ipsilateral axons was measured as a ratio of ipsilateral pixels to the total number of pixels in the dLGN region.

### RNA scope

Sections were processed according the instructions of the RNAscope Fluorescent Multiplex kit (ACDbio). No modifications were made. The RNAscope probes (ACD bio) used were: Mm-Fosb-C1, Mm-Fos-C3, Mm-Jun-C2. Fluorescent imaging was performed on a W1-Yokogawa spinning disk confocal microscope, at a 100x magnification using silicone oil. Images for Z-stacks were obtained at 0.15μm step size.

### Behavioral paradigms

For the visual cliff task, the apparatus consisted of a 60 × 30 × 30cm open-top platform with a clear bottom. The “ shallow side” laid on a flat surface and displayed directly a 4.45 × 4.45 cm black and white grid. The “ deep side” of the platform extended outside the flat surface and displayed the same grid at 2.5ft depth. Each mouse was placed on a 5×5×5 stage in the center of the platform and, behaving freely, could step onto either side. A camera was placed above the apparatus. Mice performed 10 trials within 3 days. Every trial lasted until the mouse stepped down from the stage and onto the platform, or for 5min in trials when the mice did not step down. Trials were scored as “ depth perceived” when mice stepped on the “ shallow side” of the platform, or as “ depth not perceived” when the mice either stepped on the “ deep side” or did not step down after 5min (Lim et al., 2016). Three albino mice did not step down in all 10 trials and were excluded from our analysis. For the relative size task, the apparatus consisted of the same open-top platform, with both sides laying on the same flat surface. One half of the platform displayed directly a 4.45 × 4.45 cm black and white grid, whereas the other half displayed a 1.9 × 1.9 cm black and white grid. Mice performed 5 trials within 1 day. Every trial lasted until the mouse stepped down from the stage and onto the platform, or for 5min in trials when the mice did not step down. Trials were scored as “ large-squares chosen” when mice stepped on the low spatial frequency grid, or as “ large-squares not chosen” when the mice stepped on the high spatial frequency grid. The three albino mice that did not step down in all 10 trials of the visual cliff were excluded from the relative size task as well. In both the visual cliff and the relative size task, the orientation of the platform was changed randomly between trials, to exclude behavioral influences by environmental cues.

For measurements of contrast sensitivity and visual acuity, we used the OptoDrum (Stria.Tech). Awake mice were placed on the platform inside the arena, and the OptoDrum will presented rotating stripe patterns to the animal. Stimulus rotation was set at 12°/sec. For visual acuity measurements, stimulus contrast was set at 99.72% and its spatial frequency drifted across trials from 0.061 cycles/° to 0.500 cycles/°. For contrast sensitivity measurements, stimulus spatial frequency was set at 0.061cycles/° and its contrast across trials dropped from 99.72% to 0.84%. Mouse behavior was monitored by a camera.

### Statistical analysis

Statistical analyses were performed using the Graph Pad Prism9 software. Results are presented as mean ± SEM. In all graphs, points correspond to individual mice. Two-sided unpaired t test, and two-way ANOVA (with Sidak test applied for multiple comparisons) were used with statistical significance set at p < 0.05 (*p < 0.05, **p < 0.01, ***p < 0.001, ****p < 0.0001).

---

**Figure S1: Related to Figure 1.**
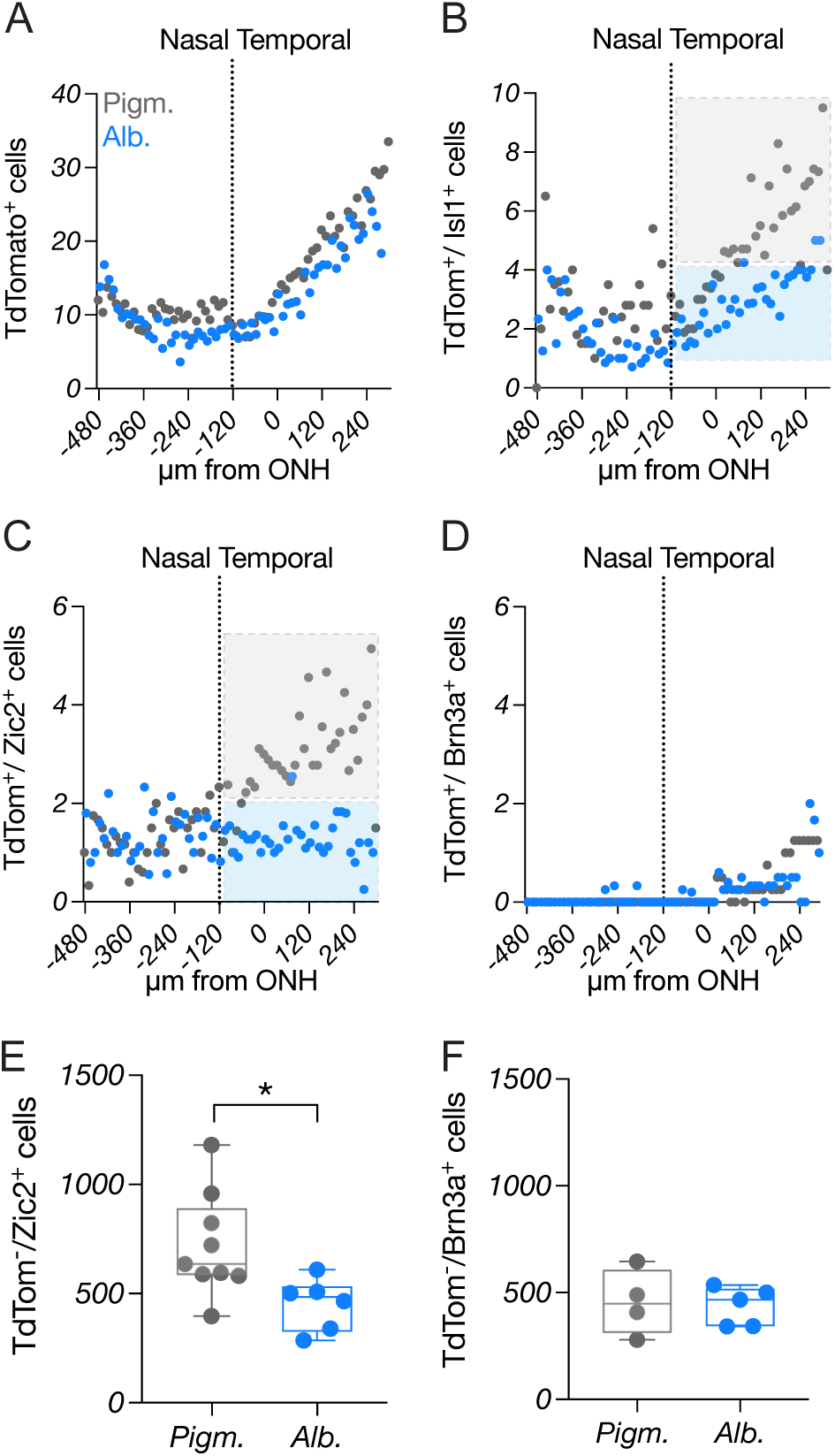
A-D) Number of tdTomato^+^ (A), tdTomato^+^/Islet1^+^ (B), tdTomato^+^/Zic2^+^ (C) and tdTomato^+^/Brn3a^+^ (D) cells in a full series of 12μm sections collected along the rostro-caudal head axis (nasal, rostral; temporal, caudal). Each point corresponds to an individual section, plotted on the x-axis as a function of distance (in μm) from the first section in the series that contained the optic nerve head opening. Note the gradual increase in tdTomato^+^ cells in the ventrotemporal retina in (A), as well as the increase in tdTomato^+^/Islet1^+^ and tdTomato^+^/Zic2^+^ cells in the ventrotemporal pigmented but not albino retina (B, C). E-F) Quantification of tdTomato^-^/Zic2^+^ (E) and tdTomato^-^/Brn3a^+^ cells in the ventrotemporal retina of pigmented and albino mice at E17.5.

**Figure S2: Related to Figure 2.**
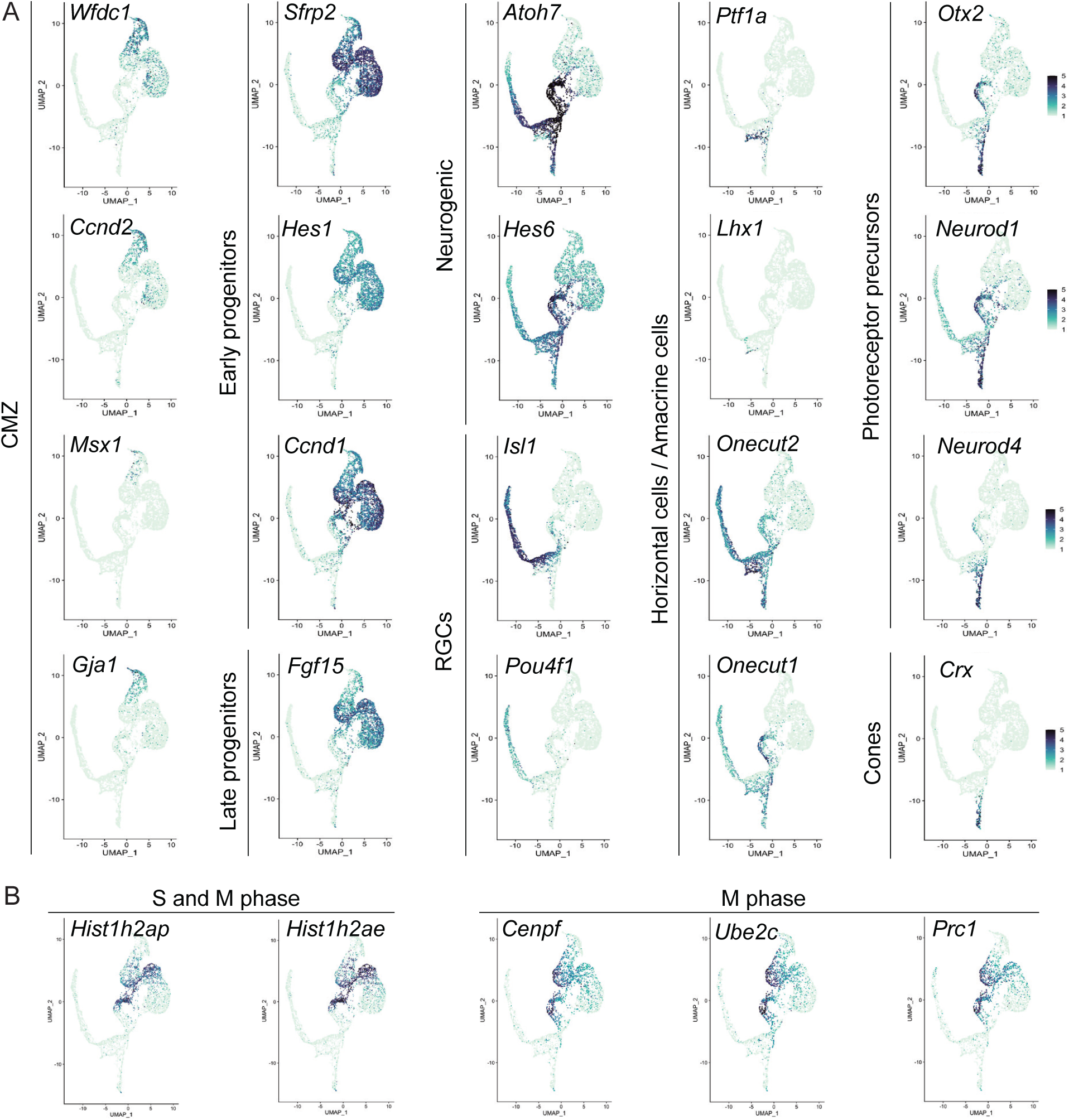
Identification of single-cell clusters by known cell type-specific markers. A) CMZ cells express *Wfdc1, Ccnd2, Msx1* and *Gja1*. Early RPCs express *Sfrp2, Hes1* and *Ccnd1* while late RPCs express *Fgf15*. Neurogenic cells express *Atoh7* and *Hes6*. RGCs express *Isl1* and *Pou4f1*. The HC/AC cluster expresses *Ptf1a, Lhx1*, and *Onecut1/2*. PR precursors express *Otx1, Neurod1* and *Neurod4*, and cones express *Crx*. B) RPC cells were enriched in *Hist1h2ap* and *Hist1h2ae*, marking the S and M cell cycle phases, as well as in *Cenpf, Ube2c* and *Prc1* marking the M phase.

**Figure S3: Related to Figure 2.**
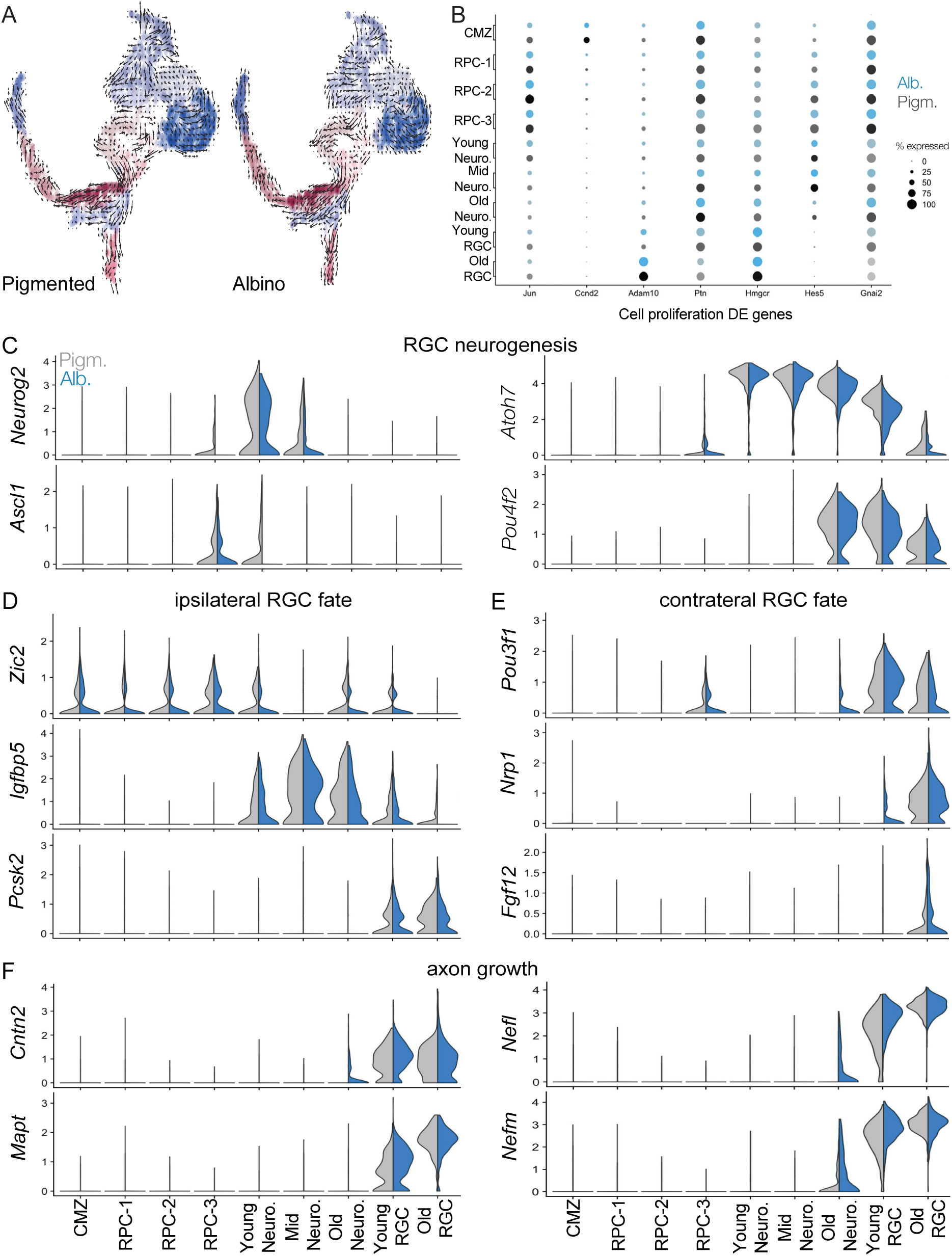
A) RNA Velocity vector fields projected on UMAP of single-cell clusters in the pigmented and albino datasets. B) Differentially expressed (DE) genes between pigmented and albino, included in the GO term “ cell proliferation” across single-cell clusters. Size of the dot represents percentage of cells within a class, while color represents average expression level across all cells within a class. C-F) Violin plots showing comparison of genes characterizing RGC neurogenesis (C), ipsilateral RGC fate (D), contralateral fate (E) and axon growth (F) between pigmented and albino scRNA-Seq datasets, across all single-cell clusters. Genes for ipsilateral RGC fate specification (*Zic2, Igfbp5, Pcsk2*) are downregulated in the albino transcriptome (D), whereas genes for contralateral RGC fate specification (*Pou3f1, Nrp1, Fgf12*) are upregulated (E).

**Figure S4: Related to Figure 4.**
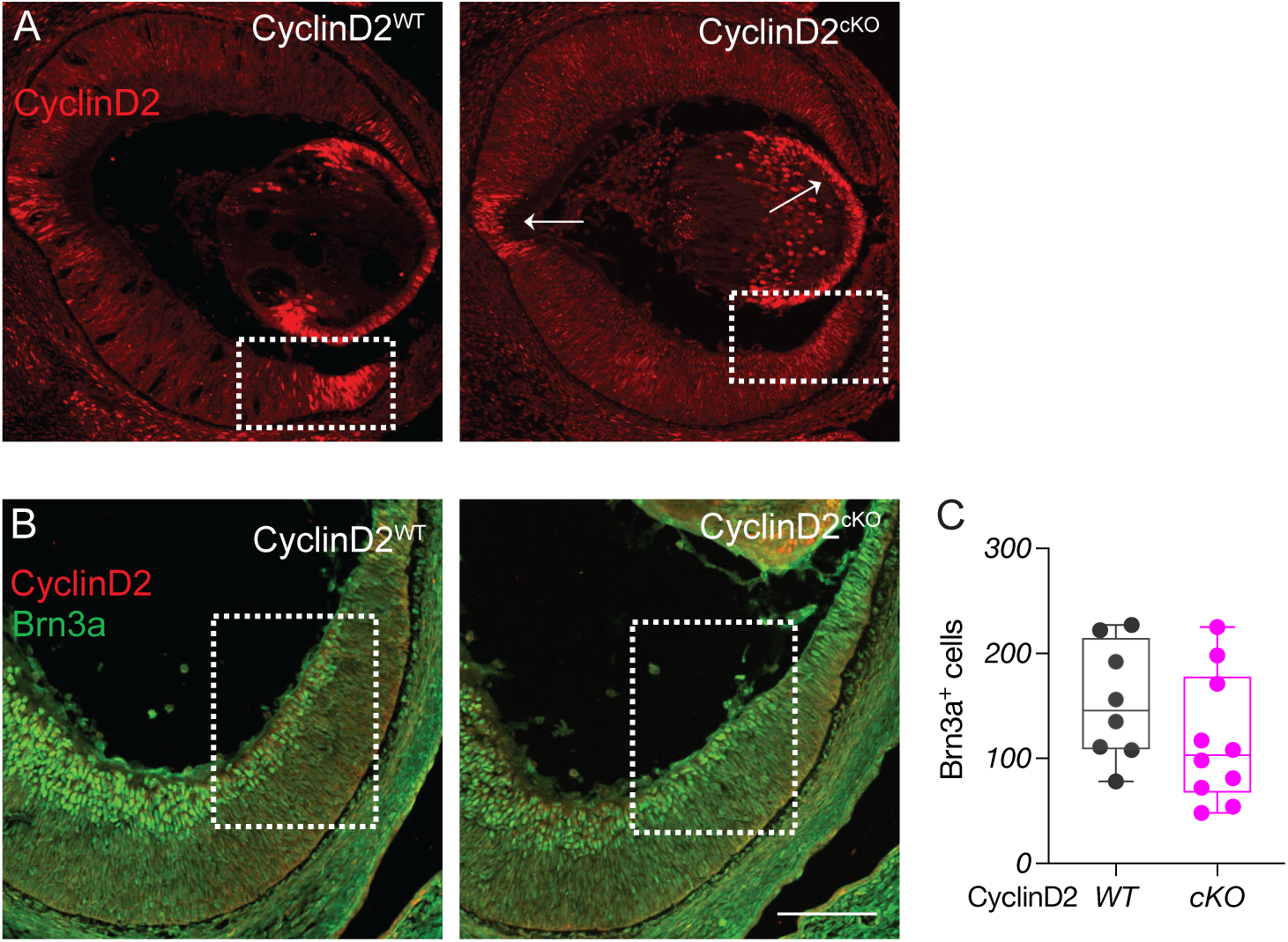
A) Example immunostaining of CyclinD2 in the CyclinD2^WT^ and CyclinD2^cKO^ ventrotemporal CMZ at E14.5. CyclinD2 is conditionally deleted from the CyclinD2^cKO^ CMZ, whereas it is still expressed in the lens and optic nerve head (arrows). B) Immunostaining of CyclinD2 and Brn3a in the CyclinD2^WT^ and CyclinD2^cKO^ ventrotemporal retina and CMZ at E15.5. Scale bar: 100μm. F) Quantification of Brn3a+ cells at E15.5.

**Figure S5: Related to Figure 5.**
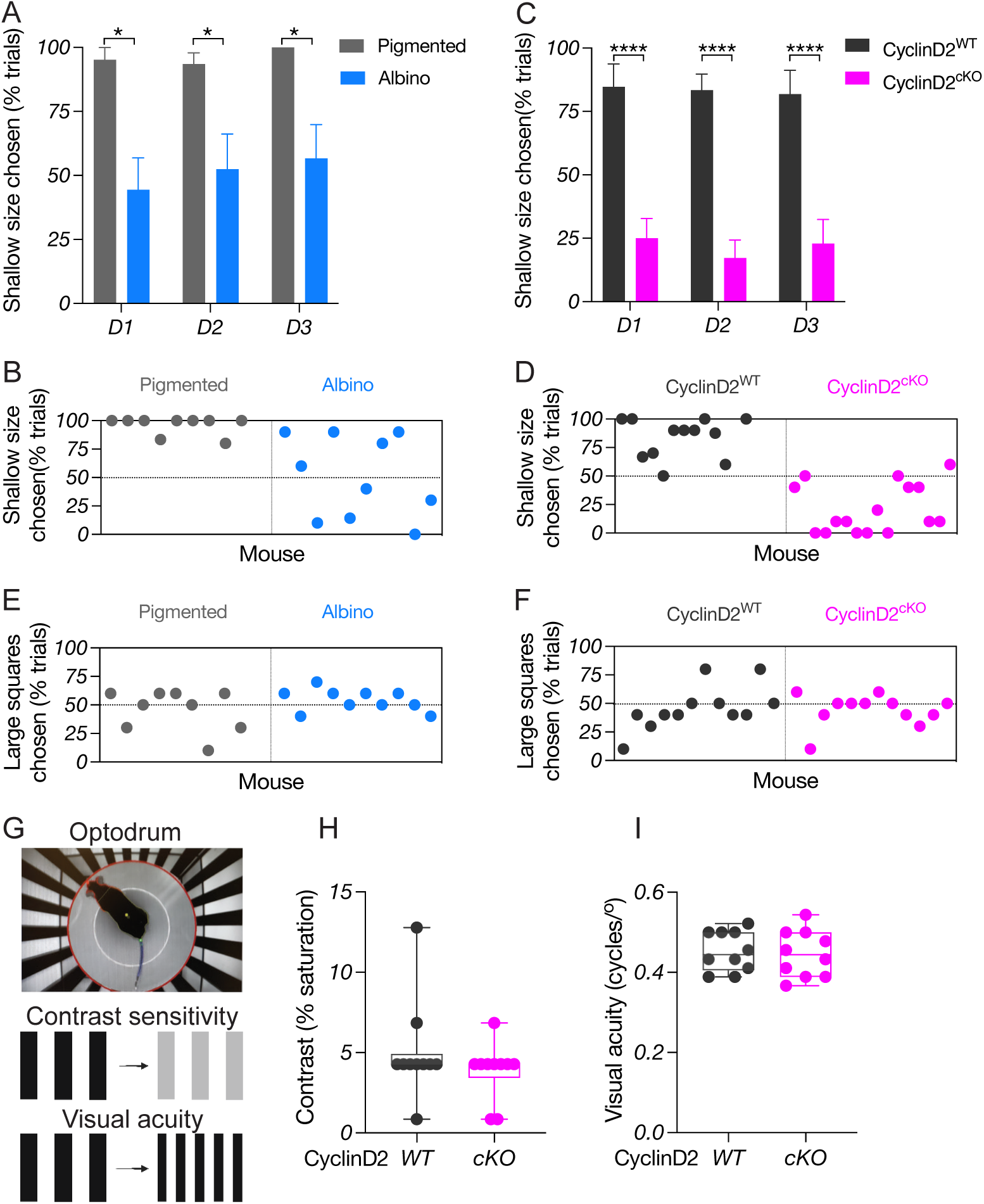
A) Quantification of depth perception at each day of the experiment, comparing pigmented and albino mice. B) Depth perception, corresponding to the percentage of trials in which the shallow side was chosen by each mouse across all trials in the visual cliff task, comparing pigmented and albino mice. C) Quantification of depth perception at each day of the experiment, comparing CyclinD2^WT^ and CyclinD2^cKO^ mice. D) Depth perception, corresponding to the percentage of trials in which the shallow side was chosen by each mouse across all trials in the visual cliff task, comparing CyclinD2^WT^ and CyclinD2^cKO^ mice. E-F) Percentage of trials in which the large-square side was chosen by each mouse across all trials in the relative size task, comparing pigmented and albino (E), and CyclinD2^WT^ and CyclinD2^cKO^ (F) mice. G) Schema of the Optodrum setup, used to measure contrast sensitivity and visual acuity. H-I) Quantification of contrast sensitivity (H) and visual acuity (I) of CyclinD2^WT^ and CyclinD2^cKO^ mice.

**Figure S6: Related to Figure 6.**
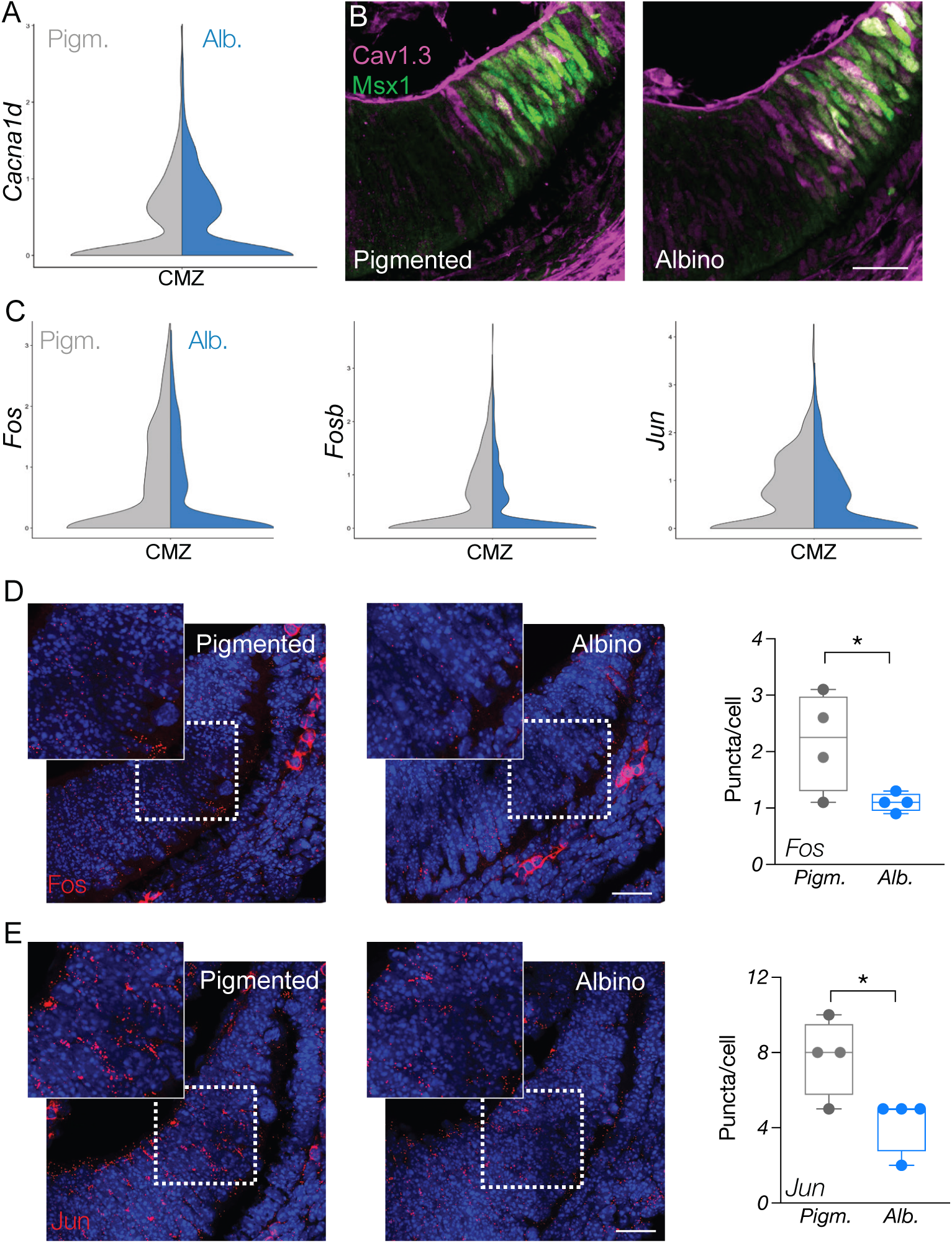
A) Violin plot of *Cacna1d* in the pigmented and albino CMZ cluster. B) Immunostaining of Msx1 and Cav1.3 in the pigmented and albino CMZ, showing Cav1.3 expression within the Msx1^+^ domain. Scale bar: 25μm. C) Violin plot of *Fos, Fosb*, and *Jun* in the CMZ cluster, showing their downregulation in the albino dataset. D-E) Fluorescent *in situ* hybridization (RNAscope) of *Fos* (D) and *Jun* (E) in the CMZ of pigmented and albino mice at E13.5. *Fos* and *Jun* expression are quantified as number of fluorescent puncta per nucleus. Scale bar: 25μm.

**Figure S7: Related to Figure 6.**
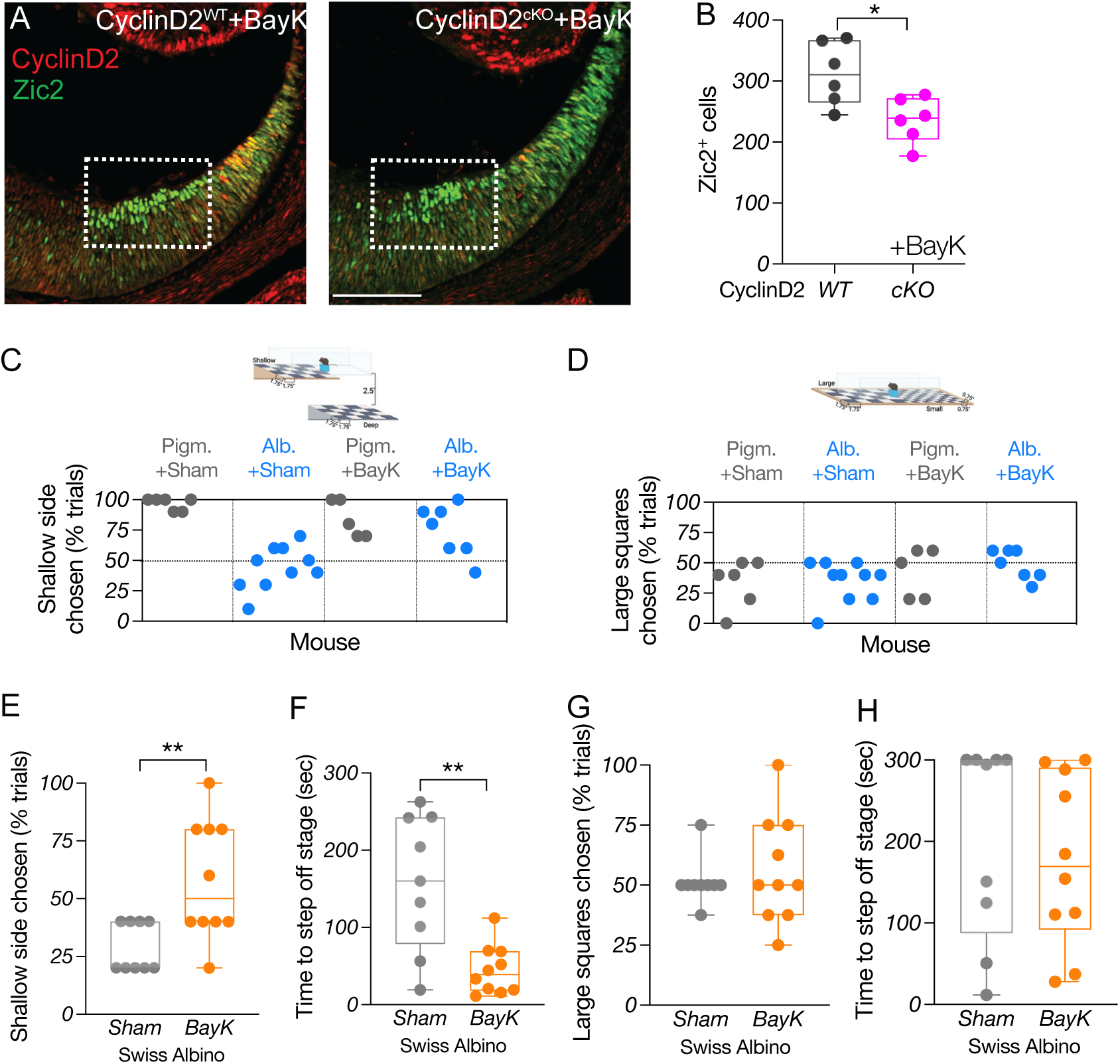
A) Immunostaining of CyclinD2 and Zic2 in the CyclinD2^WT^ and CyclinD2^cKO^, BayK-treated ventrotemporal retina and CMZ at E15.5. Scale bar: 100μm. B) Quantification of Zic2^+^ cells at E15.5. C) Depth perception, corresponding to the percentage of trials in which the shallow side was chosen by each mouse across all trials, comparing pigmented and albino Sham vs BayK treated mice. D) Percentage of trials in which the large-square side was chosen by each mouse across all trials in the relative size task, comparing pigmented and albino Sham vs BayK treated mice. E) Quantification of depth perception, corresponding to the percentage of trials in which the shallow side was chosen, comparing Swiss albino Sham vs BayK treated mice. F) Quantification of time (sec) needed to step off the stage to either side in the visual cliff task, comparing Swiss albino Sham vs BayK treated mice. G) Percentage of trials in which the large square-side was chosen, comparing Swiss albino Sham vs BayK treated mice. H) Quantification of time (sec) needed to step off the stage to either side in the relative size task, comparing Swiss albino Sham vs BayK treated mice.

